# COSMOS: integrated vector-pathogen metagenomics for mosquito-borne disease

**DOI:** 10.64898/2026.11.03.763828

**Authors:** Annie J Forster, Adil A Khan, Emma Garrett, Cheryl Toi, Carl JE Suster, John Haniotis, Kai Dang, Jack Watson, Stephen Doggett, Sharon C-A Chen, Matthew R Watts, Jen Kok, Vitali Sintchenko, Michael Payne, NSW Arbovirus Surveillance and Mosquito Monitoring Program (NSW ASMMP), Cameron Webb, Rebecca J Rockett, Tanya Golubchik

## Abstract

By 2050, almost half of the global population will be at risk of mosquito-borne disease. Effective response requires monitoring both pathogen transmission and vector abundance, yet these are typically assessed using separate assays with limited taxonomic breadth. Here we present COSMOS (COmprehensive Sequencing for MOSquito-borne disease), a targeted metagenomics workflow designed to reconstruct pathogen genomes and mosquito community composition directly from mixed field-collected samples. Over three annual rounds of a major Australian surveillance program, we evaluated >120,000 mosquitoes in 219 traps, with 98.7% agreement with surveillance data across seven monitored arboviruses, and additional detections of Sindbis virus and the state’s first detection of encephalitic Little Sussex virus. Of 137 viral detections, 101 yielded near-complete genomes (>90% coverage) at median 95x depth. Using an accompanying computational workflow, COSMOS quantified mosquito composition, matching entomological estimates of reported vector species (CCC 0.78–0.89). Joint analysis of virus and vector composition identified structured mosquito communities associated with spatial and temporal environmental variation, allowing the ecological context of arbovirus detections to be characterised. Detections during the Murray Valley encephalitis virus outbreak in 2023 were concentrated within a narrowly defined mosquito-community space corresponding to rural communities dominated by the inland S-AUS sub-lineage of *Culex annulirostris*. Integrated vector-pathogen metagenomics provides ecological and evolutionary context for arbovirus activity within a single scalable workflow for surveillance and outbreak response.

## Introduction

One in every six cases of infectious disease globally is transmitted by insect vectors, including an estimated 249 million malaria and 96 million dengue cases^1^ each year spread by mosquitoes. This burden is growing, with projections estimating that by 2050 at least 49% of the world’s population could be at risk of mosquito-borne disease outbreaks^2–4^. Emerging risk is shaped by interactions between environmental changes, vector ecology and pathogen evolution. For example, the expansion of *Aedes albopictus* into temperate regions has increased both the frequency and scale of dengue and chikungunya outbreaks^5^, compounded by emergence of viral variants with enhanced transmission by this vector^6^. Invasive mosquitoes are only part of the threat. Widely distributed *Culex* species feed opportunistically on both avian and mammalian hosts, facilitating spillover of zoonotic pathogens such as West Nile and Japanese Encephalitis viruses into human populations ^7,8^, while genetic variation among circulating viruses can further modulate disease severity^9^.

Globally, mosquito-borne disease surveillance relies on complementary but often disconnected data streams generated using different methods and at different levels of sensitivity and resolution. These include serological monitoring in humans or sentinel animals, mosquito trapping followed by expert morphological identification, and PCR-based molecular detection of a small set of expected pathogens. Although detection-based methods can be effective, they provide limited information on the origins, dispersal and persistence of viral variants or vector populations. Genomic sequencing is therefore increasingly being incorporated into arbovirus surveillance^10^, including through metagenomic^11–14^ and amplicon-based^15^ approaches. However, shotgun metagenomics may lack sufficient sensitivity for genome reconstruction^16^, whereas amplicon-based sequencing assays are challenging to scale to many species, and become vulnerable to PCR target failure as new pathogens emerge and lineages evolve^17^.

Identifying mosquito vector species presents an additional challenge. Morphological identification requires specialist entomological expertise and access to intact mosquito specimens, is highly labour-intensive, and may not distinguish closely related or cryptic taxa^16,18,19^. The handling time needed to speciate mosquitoes also contributes to RNA degradation, hampering efforts to obtain reliable viral sequence data from the same specimens. Consequently, generating linked information about vector and pathogen genomic diversity from the same specimens remains a challenge.

An integrated surveillance approach should recover pathogen genomes and mosquito community composition directly from pooled, mixed-species mosquito collections, at a scale compatible with public health surveillance. Such an approach needs to be based on a stable workflow that can identify and sequence genomes of many potentially relevant pathogens and mosquito species, while remaining robust to ongoing pathogen evolution. Targeted metagenomic sequencing using hybridisation capture has been used to recover complete viral genomes of multiple species from diverse sample types^20–22^. Separately, capture-based sequencing has also been used to resolve mosquito population structure from individual mosquitoes^23^. Importantly for surveillance applications, probe hybridisation is relatively tolerant to sequence divergence^24^, reducing the need for repeated assay optimisation as new viral lineages emerge and diversify. A combined capture panel targeting both mosquitoes and their pathogens could therefore overcome several limitations of current surveillance workflows while providing genomic resolution.

Australia provides a highly appropriate setting to evaluate this approach. Recent years have seen outbreaks of mosquito-borne disease caused by diverse endemic and emerging exotic arboviruses. These include large outbreaks of Murray Valley encephalitis virus (MVEV) in 2022/3, Ross River virus (RRV) in 2024/5, and an unexpected 2022 emergence and establishment of Japanese Encephalitis virus (JEV)^25,26^. The potential for emergence of other globally important arboviruses, such as dengue with 1,500 annual notifications largely associated with travel to Southeast Asia^27^, remains an ongoing threat. In New South Wales (NSW), Australia’s most populous state, the Arbovirus Surveillance and Mosquito Monitoring Program (ASMMP) has been operating since 1984 and collects mosquitoes every season at over 100 trap locations across a land area of 800,000km^2^, an area larger than the UK and France combined. Surveillance extends from temperate coastline, where 84% of the state’s population resides^28^, across the Great Dividing Range^29^ to sparsely populated arid inland regions. This geographic coverage encompasses diverse environments and climatic zones, which support species-rich mosquito communities, including multiple arbovirus vectors. Combined with an established virus testing program and extensive entomological expertise, ASMMP provides an ideal framework for developing an integrated targeted metagenomics approach, and evaluating its performance directly in a real-world setting.

Here, we developed, optimised and evaluated COSMOS (COmprehensive Sequencing for MOSquito-borne disease), a custom targeted metagenomics workflow designed to generate joint vector and pathogen genomic data in a single assay, combining pathogen detection, viral whole-genome reconstruction, and estimates of mosquito species diversity and relative abundance. We first benchmarked COSMOS against expert mosquito identification and routine viral testing data. We then applied the workflow directly to ASMMP mosquito collections to assess the drivers of mosquito community composition and place arbovirus detections within this ecological context.

## Results

### An integrated genomics workflow for large-scale mosquito surveillance

COSMOS was designed as a high-throughput targeted metagenomics workflow that reconstructs pathogen genomes and jointly estimates mosquito species composition directly on field-collected, mixed-species mosquito pools, without manual sorting. The workflow is summarised in **Figure 1**. Full details of method development, validation and benchmarking using *in silico* and empirical datasets are described in the **Supplementary Information**.

**Figure 1.**
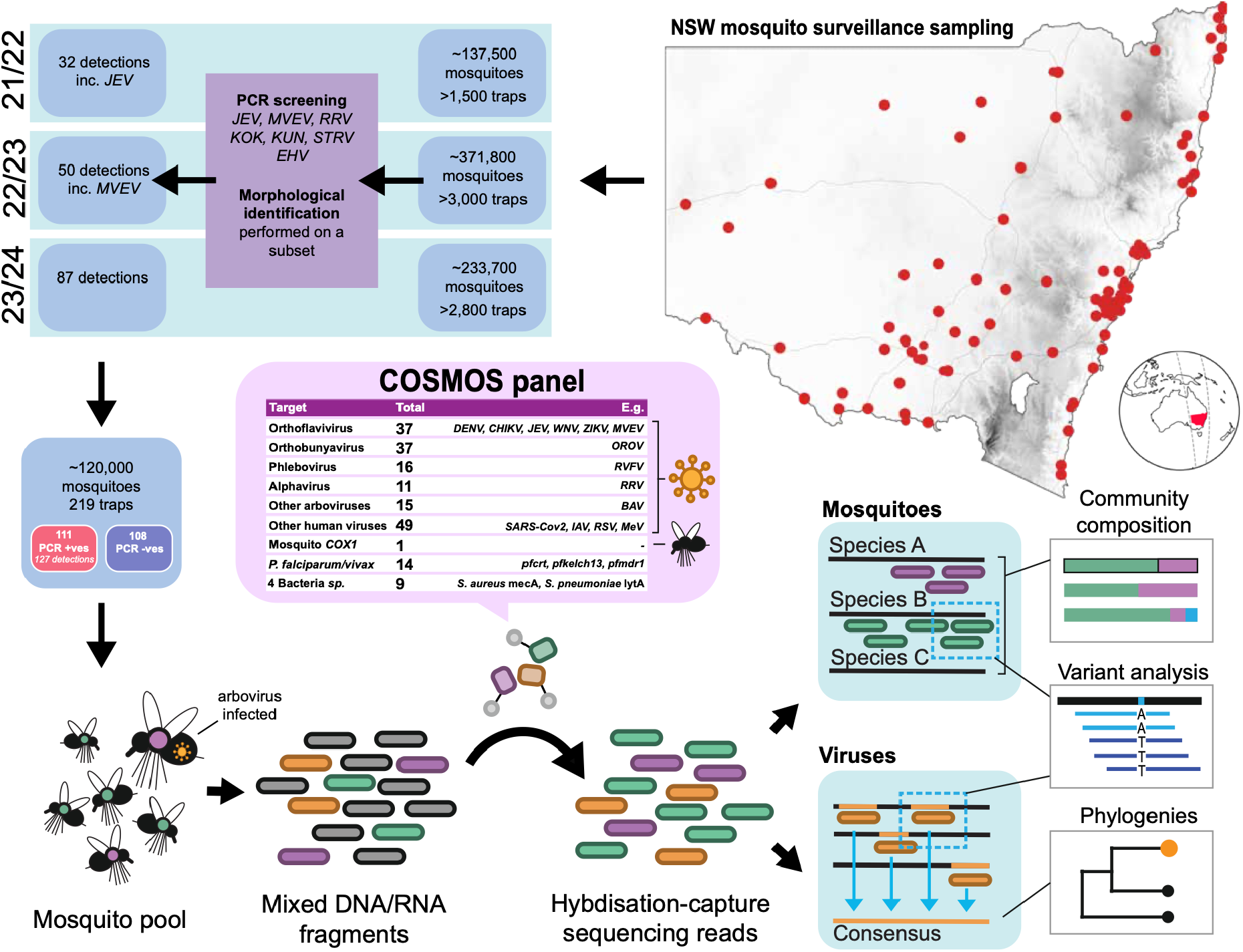
The COSMOS workflow. Map of the state of NSW shows sampling locations of surveillance traps collected by NSW ASMMP in 2021/22, 2022/23 and 2023/24 mosquito seasons (red circles). Map shading indicates altitude above sea level. Approximate numbers of trap collections and the numbers of mosquitoes they contained within these seasons are indicated, based on program records. Surveillance samples undergo routine PCR testing for the indicated arboviruses, with morphological identification performed on a subset of mosquitoes from selected traps. We prioritised 219 surveillance samples (111 PCR-positive) for the development and application of the COSMOS, an integrated hybridisation-capture genomics workflow for simultaneous recovery of virus and mosquito genetic information. *Virus acronyms: DENV: Dengue virus; JEV, Japanese encephalitis virus; WNV: West Nile virus; ZIKV; Zika virus; MVEV, Murray Valley encephalitis virus; OROV; Oropouche virus; RVFV; Rift Valley Fever virus; RRV; Ross River virus; BAV; Banna virus; SARS-CoV-2; Severe Acute Respiratory Syndrome Coronavirus 2; IAV; Influenza A virus; RSV; Respiratory Syncytial virus; MeV; Measles virus; CHIKV; Chikungunya virus; KUN; WNV Kunjin subtype; KOK; Kokobera virus; STRV; Stratford virus; EHV; Edgehill virus*.

At the completion of each of the November-April surveillance seasons in 2021/22, 2022/23 and 2023/24, we applied COSMOS to all available virus-positive samples identified by the NSW Arbovirus Surveillance and Mosquito Monitoring Program (ASMMP), together with spatially- and temporally-matched traps with no pathogen detected on routine testing. We analysed identical nucleic acid extracts from 219 traps containing >120,000 mosquitoes (111 positive by routine qPCR for at least one virus, 51%). ASMMP surveillance tests for a defined subset of locally circulating viruses and uses expert morphological identification of representative subsamples to estimate the abundance of two major vector species. We used these data to evaluate COSMOS against existing gold-standard pathogen detection and mosquito abundance estimates, while simultaneously screening for any additional pathogens and characterising mosquito community composition for the full contents of all traps.

The COSMOS hybridisation panel was designed to target the known genetic diversity of >115 arboviruses, defined as viruses with both vertebrate and invertebrate hosts, and prioritising those known to be mosquito-associated and human pathogens. The panel also included pre-validated clinically relevant loci from *Plasmodium falciparum* and their homologous regions within *P. vivax*^30^, together with 49 viral and 9 bacterial human pathogen loci to support differentiation of non-arbovirus infections and future deployment of COSMOS on clinical samples. To enable the identification of mosquito species, COSMOS targets the mosquito mitochondrial cytochrome oxidase subunit 1 (COX1) gene, a well-established genetic marker for mosquito species identification^31^, and the only gene that has been sequenced for the majority of Australian native mosquito species. Since hybridisation capture is known to be maximally efficient when probe-target sequence identity exceeds 80% ^24^, we designed probes to maintain over 80% sequence identity across the known genomic diversity of included targets, including variant arbovirus lineages, and COX1 sequences across the Culicidae family (including *Aedes, Culex* and *Anopheles* mosquitoes, as well as less-common genera such as *Uranotaenia*).

The final design comprised 48,043 probes, placing it in the second-smallest tier for custom probe design, based on quotes from two manufacturers (Twist Bioscience and Agilent). In our setting, the reagent cost for library prep, capture and sequencing was comparable to previously reported protocols for viral targeted genomics in low-resource settings ^32^, with reagent costs at AU$50 (US$35/GB£26) per sample for complete vector and pathogen assay data.

The COSMOS panel design and target composition are listed in **Table S1**. For full reproducibility, we have made the panel available as a defined custom reagent (this will be available through Twist Bioscience as design ID TE_XXXX).

### COSMOS simultaneously recovers viral genomes and mosquito genetic information

COSMOS efficiently enriched both mosquito and viral genomic targets. High enrichment was observed under both controlled laboratory conditions using pre-defined mosquito pools (**Fig. S1A** and detailed in **Supplementary Information**), and on field-collected mosquito surveillance samples. Across all surveillance samples, median depth was 1,215x (IQR 240–4,156) at the COX1 control target, with approximately 471x enrichment relative to the untargeted fraction of the mitochondrial genome (**Fig. 2A**). Similarly, viral genomes were recovered at high depth (median 95x, IQR 11–395) and coverage (median 99.3% complete, IQR 86.5–100%) across all 137 COSMOS detections (**Fig 2B**), and 101/137 (73%) genomes were near complete (>90% genome coverage). COSMOS results were overall 98.7% concordant with ASMMP qPCR results, with 90% of qPCR-positive detections yielding a positive COSMOS detection (N=114/127), and an additional 10 detections from COSMOS that had been missed by qPCR. Genome recovery by COSMOS was proportional to input viral load, consistent with successful enrichment: 50% genome coverage was predicted at a qPCR cycle threshold (Ct) of 27.7 (26 - 29.38 95% CI) and 95% coverage at Ct 26.1 (21.67 – 27.07 95% CI), based on a logistic model fitted to the observed data (**Fig. 2C**).

**Figure 2.**
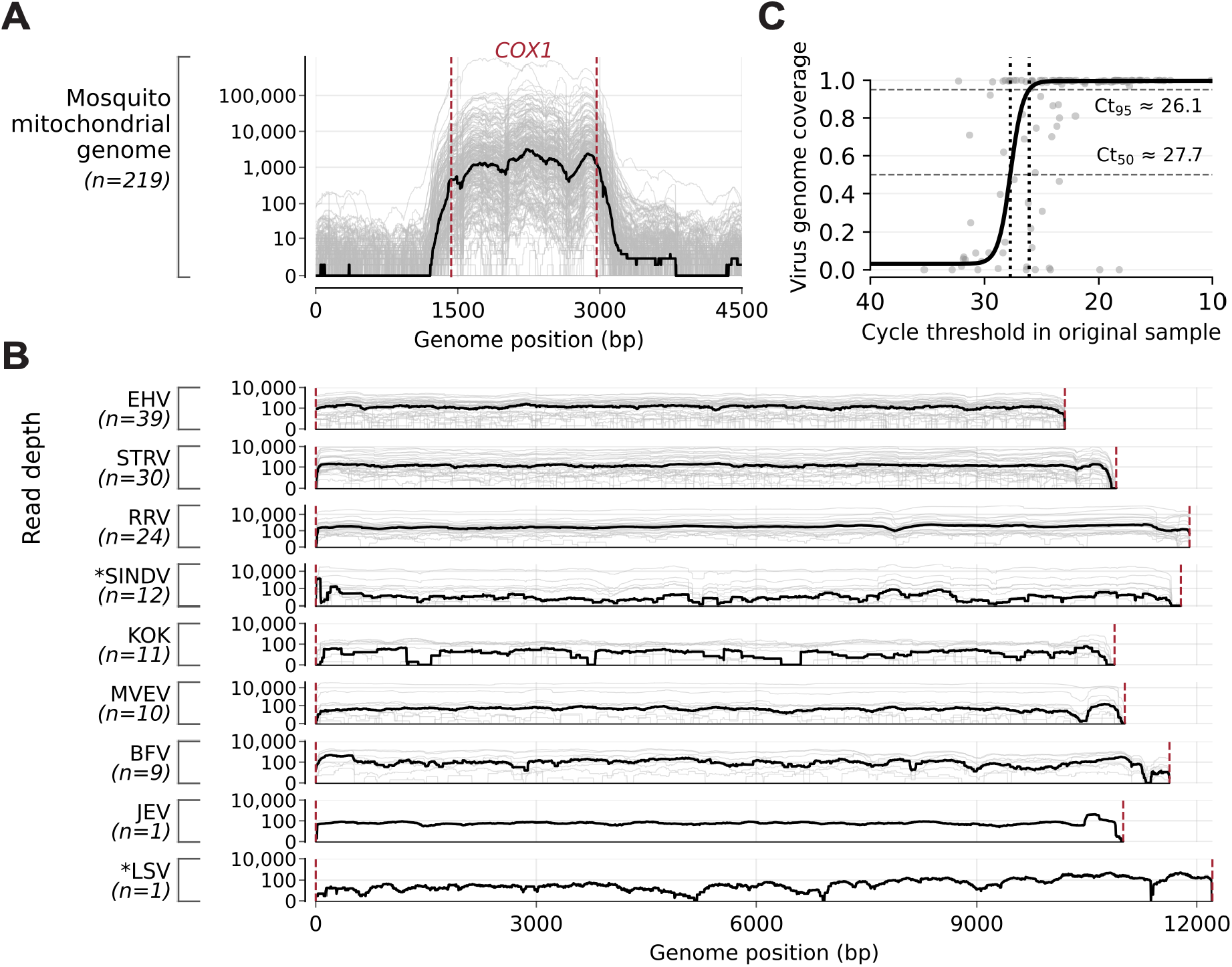
COSMOS enriches mosquito and viral genetic targets from pooled surveillance samples. Enrichment at **(A)** the mosquito *COX1* gene on the *Culex annulirostris* mitochondrial genome and **(B)** viral genomes detected by COSMOS. Median depth (thick black line) and individual sample depth (fine gray lines) shown for all field-collected samples. Red dashed lines indicate boundaries of targeted regions (whole genomes for viruses). Asterisk next to LSV and SINDV indicates new COSMOS detections. (**C**) Comparison of genome coverage vs reported Ct values for viruses targeted by both ASMMP surveillance qPCR and COSMOS sequencing. The fitted four-parameter logistic curve is shown, with Ct_50_ and Ct_95_ indicating the Ct values at which the model predicts 50% and 95% genome coverage, respectively. *Virus name abbreviations: EHV, Edge Hill virus; STRV, Stratford virus; RRV, Ross River virus; SINDV, Sindbis virus; KOK, Kokobera virus; MVEV, Murray Valley encephalitis virus; BFV, Barmah Forest virus; JEV, Japanese encephalitis virus; LSV, Little Sussex virus*.

Detections in field-collected samples belonged to nine arbovirus species, seven of which were screened by routine ASMMP surveillance: the flaviviruses JEV, MVEV, Kokobera virus, Edge Hill virus, Stratford virus, and the alphaviruses Ross River virus and Barmah Forest virus (West Nile virus Kunjin variant is also included in routine ASMMP surveillance but was not detected by ASMMP testing or COSMOS during the sampling period). In addition, COSMOS recovered a near-complete genome of Little Sussex virus (LSV), which had never previously been detected in NSW and had been reported only once before in Australia, in Queensland in 1976 ^33^. COSMOS also identified 12 surveillance samples positive for Sindbis virus across all three sampled seasons, including 5 complete genomes consistent with high viral load (**Fig. 2B**). Sindbis virus had not been included in routine PCR testing during COSMOS sequencing, and its detection has prompted subsequent inclusion in the ongoing ASMMP surveillance. Beyond this, analysis of residual metagenomic background sequence also recovered a broader mosquito-associated virome (**Fig. S2** and **Supplementary Results**).

### COSMOS estimates of mosquito community composition are concordant with expert morphological identification

A major goal of COSMOS was to use sequencing data to provide a robust and scalable alternative to morphological mosquito identification and quantification, which is labour-intensive and challenging for closely related taxa. Numerous studies have established that hybridisation-capture sequence recovery scales with input target abundance ^22,32,34^, and COSMOS showed this quantitative behaviour for viral targets: the number of unique sequencing reads was strongly correlated with viral load in laboratory-derived serial dilutions (**Fig. S3A**) and across field-collected samples (R^2^=0.47, P=8.11×10^−19^; **Fig 3A**). We therefore reasoned that it may be similarly quantitative for mosquito targets, which would enable the estimation of mosquito species relative abundance directly on mixed-species surveillance samples, provided species could be identified reliably and at comparable sensitivity.

**Figure 3.**
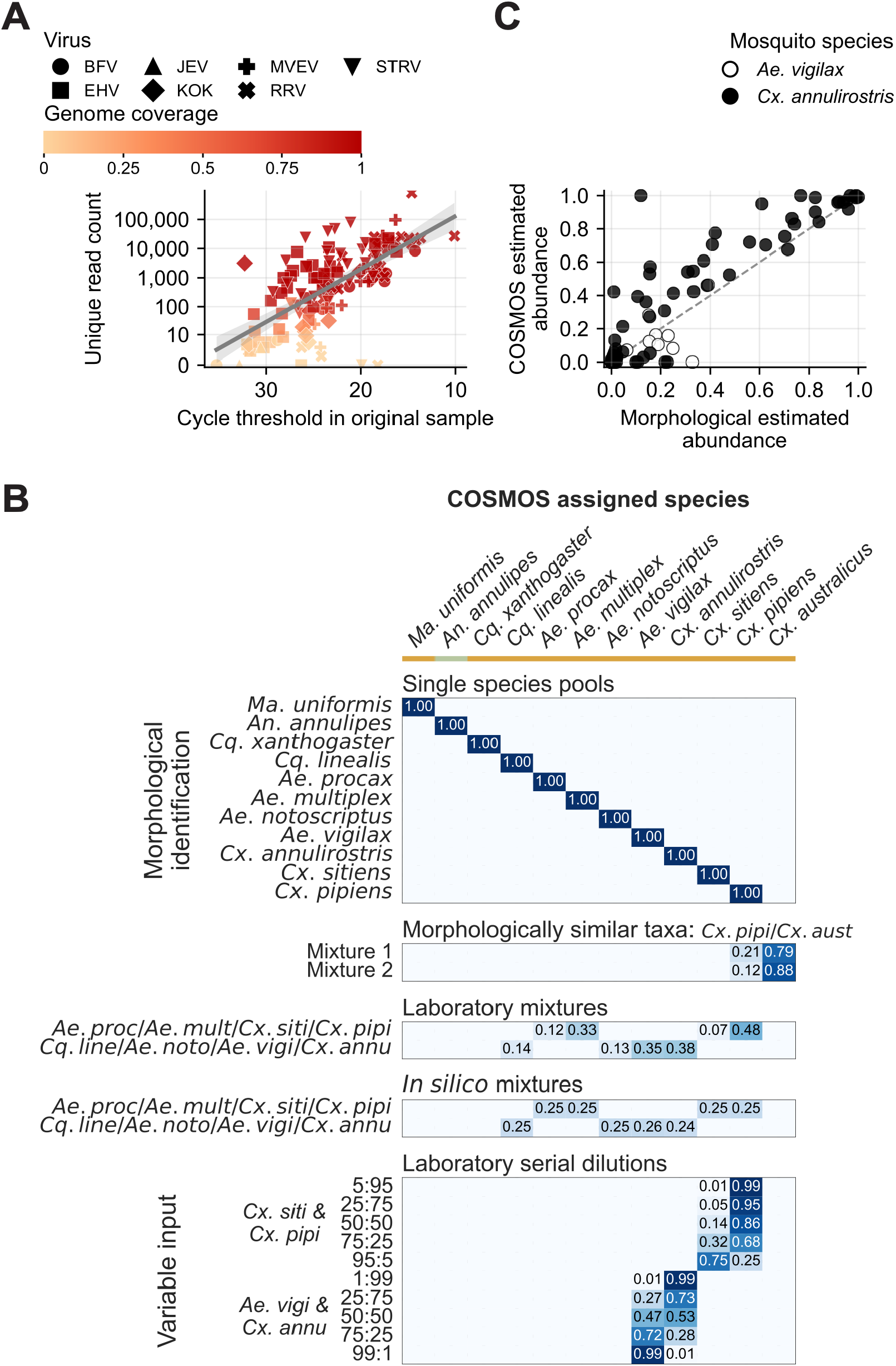
COSMOS provides quantitative viral and mosquito community information from pooled samples. **(A)** Each point shows a PCR-positive virus detection across surveillance traps (N=127). The x-axis shows the qPCR cycle threshold (Ct) for each detection, which is an inverse proxy for viral load within the sample. The y-axis shows the number of unique reads assigned to each virus by Castanet from COSMOS-enriched sequence data, shown on a log_10_ scale. Points are coloured by genome recovery category reflecting the proportion of the virus genome covered by at least two reads and are shaped by the identity of each virus. Across ASMMP detections (N=127), a linear regression was performed which fitted the model: log_10_(unique read count + 1) = 6.93 - 0.18 × Ct (R^2^=0.47, P=8.11 ×10^−19^). The fitted line is shown in grey with a shaded 95% confidence interval. **(B)** Heatmaps show estimated relative abundance of mosquito species in the reference database (x-axis) across test samples (y-axis). “Single-species pools” shows morphologically identified single-species mosquito pools enriched using COSMOS. “Morphological ID comparison” shows two pools of ten mosquitoes putatively identified morphologically as *Cx. australicus*, a species that is difficult to distinguish morphologically from members of the *Cx. pipiens* complex. “Laboratory mixtures” shows experimental mixtures containing equal numbers of individuals from four mosquito species. “Simulated mixtures” shows *in silico* mixtures generated by combining equal proportions of reads from single-species pools to match the species composition of the corresponding laboratory mixtures. “Laboratory serial dilutions” shows two serial dilution experiments combining pairs of mosquito species at varying input proportions, with expected proportions indicated on the x-axis labels. *Culex pipiens* denotes the analytical group containing *Cx. pipiens, Cx. quinquefasciatus* and other members of the *Cx. pipiens* complex that were sequence-identical across the COX1 region used for species assignment; *Cx. australicus* was distinguishable and is reported separately. *An*.= *Anopheles, Cq*.= *Coquillettidia, Ma*. = *Mansonia*. **(C)** Each point represents a surveillance trap with a detection of *Aedes vigilax* (N=72) (not filled) and *Cx. annulirostris* (N=75) (filled) and a location and date matched entry in NSW Mosquito Surveillance Programme published mosquito counts. The x-axis shows the relative abundance of each species compared with a count of total mosquitoes within the trap, and the y-axis shows the relative abundance estimated using COSMOS.

To quantify mosquito species abundance using COSMOS-derived sequence data, we developed a workflow based on probabilistic read assignment against a shared locus. The COSMOS mosquito quantification workflow was designed to use a curated reference database of 1kb mosquito COX1 sequence, representing species and/or species groups relevant to Australia and the Indo-Pacific region, as described in **Methods**. Some related members of the *Cx. pipiens* complex, including *Cx. pipiens molestus* and *Cx. quinquefasciatus*, were sequence-identical at the reference region and were therefore grouped into a single assignment category labelled ‘*Cx. pipiens*’ for simplicity. Reference sequences were obtained from all available species; for most Australian species, COX1 was the sole locus available in GenBank.

We evaluated the workflow against a set of samples of known composition based on expert morphological identification by a highly experienced entomologist familiar with Australian mosquito collections (**Fig. 3B**). On 11 single-species sorted pools of 20 individual mosquitoes each, representing the most commonly found vector species in the NSW region, all filter-passing signal was assigned to the expected species. Because *Cx. australicus* was challenging to distinguish morphologically from other members of the *Cx. pipiens* complex, we additionally tested the COSMOS workflow on two pools of ten mosquitoes each that had been morphologically characterised as *Cx. australicus*. In these pools, 20.5% and 12.2% of signal was instead assigned to the *Culex pipiens* group respectively, and further analysis showed consistent minority *Cx. pipiens* group alleles across species-informative sites, supporting the presence of a mixture of species rather than ambiguous assignment (**Fig. S4**). This highlights the value of the COSMOS approach for resolving visually similar taxa.

Next, we tested whether COX1 abundance retained information about species proportions within mixed species samples. In *in silico* mixtures generated by combining reads from four individually sequenced single-species pools, all component species were identified with minimal error overall (**Fig. S5**). In two laboratory-based mixtures containing equal numbers of four mosquito species, all four input species were again detected in both mixtures, although their estimated proportions deviated from the expected equal composition (**Fig. 3B**). This contrast suggests that variation in estimated abundance was introduced prior to computational read assignment, potentially reflecting differences in mosquito biomass, mitochondrial copy number or extraction/capture efficiency. Nevertheless, in serial dilutions of two sets of mosquito-species pairs, estimated abundance changed in the expected direction with input ratio, supporting the use of COSMOS to detect relative shifts in mosquito composition (**Fig. 3B)**.

Finally, we applied the workflow to the 219 surveillance samples and compared COSMOS-derived species proportions with routine ASMMP morphological identification estimates. These were made based on counts of *Cx. annulirostris* and *Ae. vigilax* within a subset of mosquitoes within each trap and available for a subset of traps. Concordance between COSMOS-derived and morphology-based estimates was high for *Cx. annulirostris* (Pearson’s R=0.92, Lin’s concordance correlation coefficient (CCC)=0.89, N=75) and moderate-to-high for *Ae. vigilax* (Pearson’s R=0.79, CCC=0.78, N=72) (**Fig. 3C**), indicating that COSMOS provides reliable estimates of relative mosquito species abundance in field-collected surveillance samples.

Together, these analyses support the use of COSMOS to identify mosquito species and detect relative differences in community composition in routinely collected surveillance material. Combined with simultaneous viral genome recovery, COSMOS therefore enables pathogen genomic diversity and vector community composition to be resolved together from the same samples and within a single scalable workflow.

### Identifying ecological correlates of arbovirus circulation within mosquito communities

To profile mosquito species composition and abundance across the state, we examined 213/219 (97.3%) of surveillance traps with sufficient COX1 sequence data **(Fig. 4A** and **Fig. S1B)**. Samples without estimates had low COX1 coverage (median 0.24, range 0.04–0.31) compared to those with (median 1, range 0.81–1 **Fig. S1B**). In total, 23 mosquito species were identified, with *Cx. annulirostris* the most frequently detected, present in 170 traps (**Table S2**). This species has been recognised as the dominant vector of multiple clinically important viruses, including JEV and MVEV ^35,36^.

**Figure 4.**
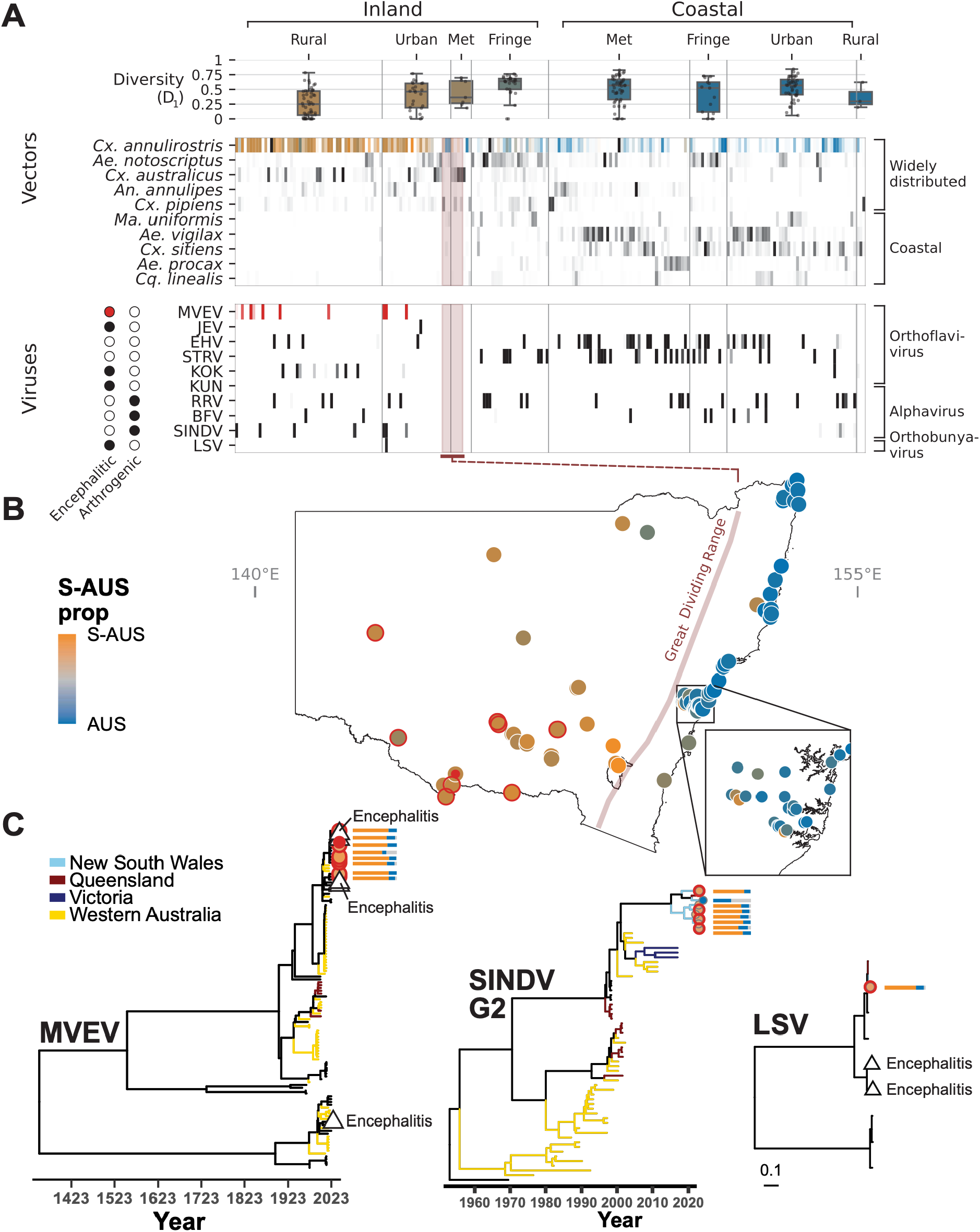
Joint vector–virus profiling contextualises arbovirus outbreaks in New South Wales. In all panels, colour indicates estimated proportion of *Cx. annulirostris* lineage S-AUS (inland, orange) and AUS (coastal, blue), and detection of MVEV is marked in red. The position of the Great Dividing Range is indicated in dark red. Panel A shows mosquito traps as coastal or inland grouped by council classification, separated by grey lines. Council groups are ordered by their mean longitude, with traps within each group ordered from west to east. The region corresponding to the Great Dividing Range is shaded to link the heatmaps with the corresponding geographic region in panel B. **(A)** Boxplots show mosquito species diversity across area classifications for inland (left; >20 km from the coast) and coastal (right; ≤20 km from the coast) surveillance traps. Each point represents a trap with mosquito species assignments (N=213), and boxplots show the distribution within each classification. Traps were classified as rural, urban, metropolitan fringe (‘Fringe’) or metropolitan (‘Met’) according to NSW council classifications, with Australian Capital Territory sites classified as metropolitan. Diversity is calculated as Simpson’s diversity index (*D*_1_), with higher values indicating greater mosquito species diversity. Each boxplot is shaded by mean S-AUS proportion within each group. The upper heatmap shows estimated relative abundance of the ten most frequently detected mosquito species while the lower heatmap shows COSMOS detected viruses in the same 219 surveillance traps (columns). Mosquito species are ordered by detection frequency and *Cx. annulirostris* is coloured by lineage proportion, from inland S-AUS (orange) through coastal (AUS) blue, with colour intensity proportional to estimated relative abundance. *Culex pipiens* denotes members of the *Cx. pipiens* complex, which are sequence-identical at the COX1 region used for species assignment. For viruses, colour indicates the proportion of the viral genome covered. **(B)** Geographic distribution of *Cx. annulirostris* COX1 lineages S-AUS (orange) and AUS (blue). Each point represents a surveillance location, coloured by lineage proportion estimated from read pile-ups at four lineage-informative sites identified from publicly available sequences reported by Hemmerter *et al*. (2007). Traps were included where >20 reads covered all four sites (N=157). Where multiple traps came from the same location, lineage proportions were averaged, resulting in 78 mapped locations. MVEV-positive traps are indicated by red-outlined circles (9 detections across 8 sites). One additional MVEV-positive trap for which mosquito community composition could not be resolved is shown as a solid red dot. **(C)** Time-resolved phylogenies of MVEV and Sindbis virus and a phylogeny of Little Sussex/Umbre viruses. COSMOS-derived sequences carrying MVEV are outlined in red across all trees. Accompanying bars show the estimated proportion of *Cx. annulirostris* within the mosquito community, subdivided according to the estimated AUS (blue) and S-AUS (orange) lineage proportions. Where a sample was collected within Australia, the branch colour indicates the state of origin, as described in the legend. One MVEV detection had insufficient genome coverage for inclusion in the phylogeny. The MVEV-positive trap lacking mosquito community information is indicated by a solid red dot.

Previous work has identified at least five deeply diverged mitochondrial COX1 lineages of *Cx. annulirostris* across the Indo-Pacific of which two, “S-AUS” and “AUS”, currently circulate in Australia ^37^. Using COSMOS-derived sequence data, we estimated the proportion of AUS and S-AUS *Cx. annulirostris* lineages using allele frequencies at lineage-defining polymorphic sites in COX1, in the 157/170 (92%) of *Cx. annulirostris*-positive traps with high read support (see **Methods**). AUS and S-AUS lineages showed strikingly distinct spatial distributions (**Fig. 4B**); the average proportion of S-AUS-type base calls increased strongly with distance from the coast (N=157, linear regression R^2^=0.54, P<0.0001**; Fig. S6**) with the transition between S-AUS and AUS lineages closely coinciding with the Great Dividing Range, a mountain range running down the southeast coast of Australia (**Fig. 4A**,**4B**). Based on mitochondrial and microsatellite analyses, the Great Dividing Range is thought to act as a barrier to gene flow between coastal and inland *Cx. annulirostris* populations ^38^, and we therefore considered these lineages separately in subsequent analyses of community composition.

We next asked which environmental and temporal factors might shape variation in mosquito community structure across NSW. We found that the first principal component (PC) was spatially driven, strongly associating with distance to coast (marginal R^2^=0.519, FDR-adjusted P=1.81 ×10^−28^) and land-use type (marginal R^2^=0.319, FDR-adjusted P=7.31 × 10^−13^), with rural areas showing the greatest effect, possessing lower species diversity (**Fig. 5** and **Fig. S7**). The second major axis (PC2) showed stronger associations with temporal variables, including season (marginal R^2^=0.105, FDR-adjusted P=6.135×10^−7^) and sampling week (marginal R^2^=0.055, FDR-adjusted P=0.0002), reflecting seasonal turnover in communities; *Culex australicus*, which typically peaks earlier in the surveillance season ^39^, contributed most strongly to this axis.

**Figure 5.**
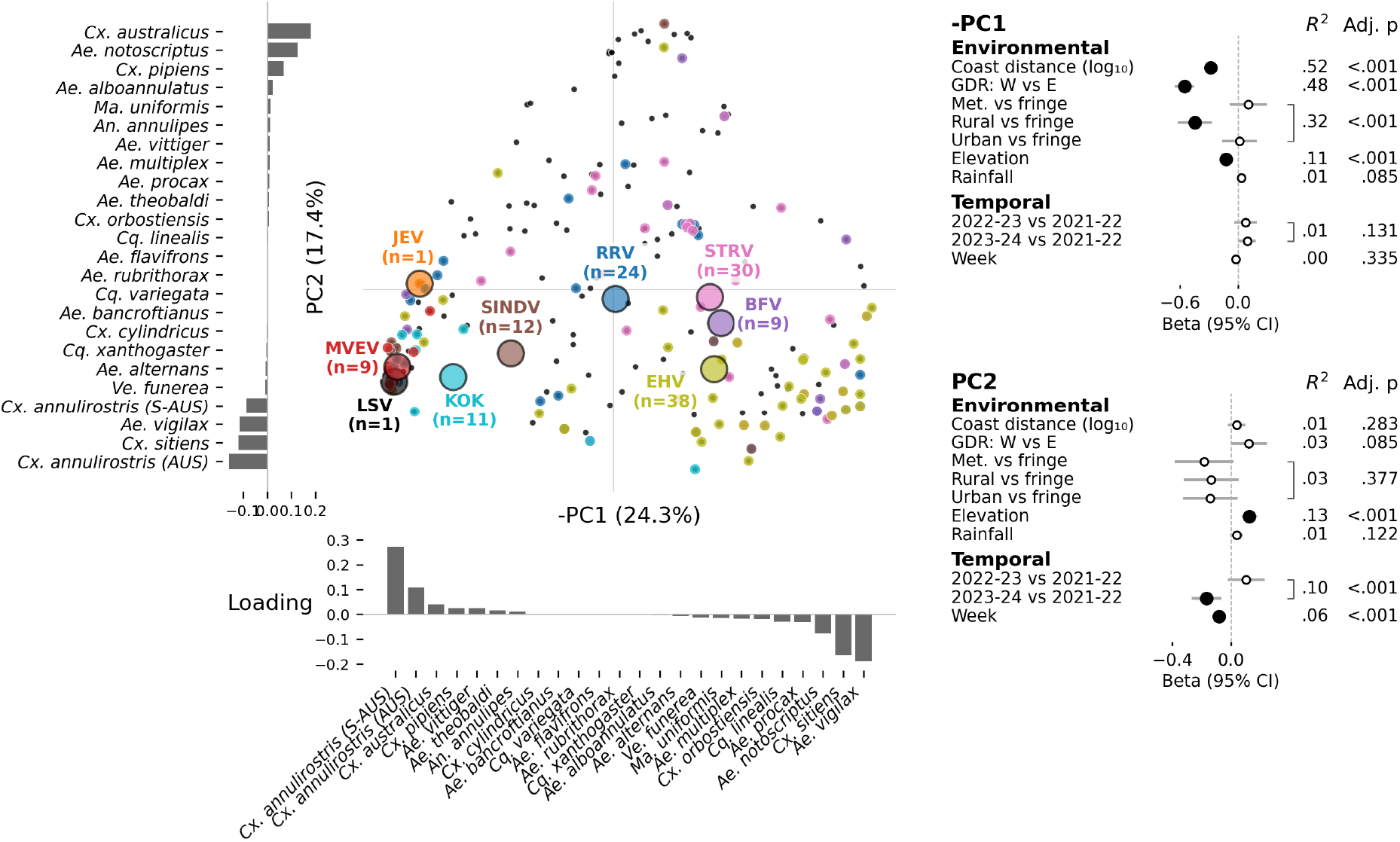
Arboviruses circulate within ecologically structured mosquito communities. Each point in the ordination represents a surveillance sample with mosquito abundance estimates (N = 213); larger coloured points indicate traps in which an arbovirus was detected. For traps containing multiple viruses, the least frequently detected virus across the dataset is shown. Principal components (PCs) were calculated from Hellinger-transformed abundances of 24 mosquito species (treating *Cx. annulirostris* S-AUS and AUS as separate; **Fig. 4**), with PC1 (x-axis) oriented to reflect the west-to-east geographic gradient and PC2 shown on the y-axis. Large, labelled points show the centroid of detections for each virus in mosquito-community space. Bar plots aligned with each axis show the loading of each mosquito species on the corresponding PC, ordered by loading. Forest plots show associations between selected environmental and temporal variables and PC1 or PC2, estimated using linear mixed models with sampling location included as a random intercept to account for repeated sampling. Points and lines show fixed-effect coefficients (β) and 95% confidence intervals; marginal R^2^ and p-values are annotated. P-values were corrected across PC-association tests using the Benjamini–Hochberg method. For categorical predictors, the displayed coefficients represent contrasts with the indicated reference category, while statistical significance of the predictor was assessed using a joint Wald test. Associations with FDR-adjusted P < 0.05 and 95% CI excluding zero are highlighted. PC1 regression coefficients were reversed together to match the displayed orientation of the PC1 axis. *Ve. = Verrallina*.

We projected arbovirus detections onto this mosquito-community space, which showed that viruses vary in the type and breadth of mosquito community contexts which they occupy (**Fig 5**). For example, the vector generalist RRV ^40^ was broadly distributed across this ordination, spanning multiple vector community states, while MVEV detections were tightly clustered within a narrower community context. These detections occurred during the unexpected summer 2023 outbreak of MVEV in southeastern Australia^26^, providing an opportunity to further examine viral genomic diversity within its identified mosquito community context.

### Cryptic vector population structure provides ecological context for the 2023 Murray Valley encephalitis virus outbreak in NSW

We therefore examined MVEV genome diversity from the 2023 outbreak. MVEV encephalitis is associated with high human mortality, with case fatality rates between 15-30% ^41^. All but one (9/10) COSMOS detections of MVEV yielded a consensus genome covering >30% of the expected MVEV genome (coverage range 31–100% with 6 >99% complete). Phylogenetic analysis combined with publicly available MVEV genomes confirmed that, like other sequences collected in other southeastern Australian states during this outbreak, these sequences belonged to the Genotype 1A^26^. Mosquito species abundance could be estimated in 9/10 MVEV-positive traps, in which *Cx. annulirostris* dominated, accounting for a median estimated mosquito abundance of 99% (range, 76–100%). Within these traps, the *Cx. annulirostris* signal was overwhelmingly assigned to the S-AUS lineage (S-AUS proportion range 0.782 – 0.872), consistent with its inland geographic distribution (**Fig. 4C**).

Notably, MVEV circulation co-occurred with other arboviruses. Four of the 12 Sindbis virus (SINDV) detections co-occurred with MVEV, and one of these surveillance samples also contained Little Sussex virus (LSV), never previously seen in NSW (**Fig. 4C**). Neither SINDV nor LSV were part of routine screening, and both were first detected by COSMOS, enabled by the platform’s broad target coverage.

In particular, among 8/12 SINDV genomes available for phylogenetic analysis we detected two co-circulating lineages. Six genomes formed one lineage detected exclusively inland, with one detection in 2021-22 and five in 2022-23. The remaining pair of genomes formed a second lineage detected inland in the 2022-23 season, co-occurring with the first lineage at the same sampling site, and once on the northern coast the following season. Within the second lineage, the coastal genome was most closely related to a publicly available sequence collected in 2022 in Queensland, which is immediately north of NSW. This pattern suggests spatial and temporal movement of SINDV within NSW, which its addition to routine surveillance and further sequencing may help to resolve (**Fig. 4C**).

Very little is known about LSV, which is closely related to Umbre virus, another understudied arbovirus. Recently, Umbre virus was identified as the cause of two fatal cases of encephalitis in France^42^ (**Fig. 4C**). Whether these patterns indicate shared ecological drivers of arbovirus activity is of distinct interest to further research.

Together, these analyses place MVEV genomic diversity within a geographically structured *Cx. annulirostris* landscape, showing it is possible to resolve viral relatedness, mosquito community composition and cryptic vector population structure from genetic data directly from the same surveillance traps.

## Discussion

Understanding mosquito-borne disease transmission and outbreak risk requires consistent information on mosquito vectors and the pathogens they carry, yet these are typically not assessed together. Even when both are collected, such as by the NSW Arbovirus Surveillance and Mosquito Monitoring Program (ASMMP) in Australia, the ability to integrate broad vector and pathogen information remains constrained by targeted pathogen testing and the availability of entomological expertise. In this study, we developed a novel targeted metagenomic workflow COSMOS (**Fig. 1**) that profiles diverse mosquito communities without requiring morphological mosquito identification, and reconstructs viral genomes at sensitivity comparable with standard PCR-based molecular assays, directly from the same samples **(Fig. 3)**. Applied to mixed-species field collections from the ASMMP, COSMOS revealed spatially and temporally structured mosquito communities, allowing the ecological context of arbovirus detections to be characterised (**Figs. 4** and **5**).

COSMOS was also capable of sub-species resolution, identifying geographically structured mitochondrial lineages in *Cx. annulirostris* and *Ae. notoscriptus* (**Fig. 4** and **Fig. S8**). *Cx. annulirostris* is a particularly important Australian vector due to its ability to spread multiple clinically significant arboviruses. It has long been associated with transmission of MVEV, and is the major vector in the ongoing spread of Japanese encephalitis virus ^43^. Across the Indo-Pacific, multiple divergent lineages of *Cx. annulirostris* have been identified using mitochondrial and genomic markers ^37,38^. We observed strongly contrasting spatial distributions of the two Australian mitochondrial COX1 lineages, AUS and S-AUS, with a marked transition across the Great Dividing Range, supporting previous inferences of a partial barrier between these subpopulations ^38^ (**Fig. 4**). Within this geographic structure, COSMOS generated the first whole genomes of MVEV from samples collected in NSW and placed the reconstructed genomes within the G1A subclade, alongside genomes sequenced from the 2023 outbreak in Victoria, a neighbouring state ^26^. The corresponding traps were dominated by *Cx. annulirostris*, predominantly of the S-AUS lineage, consistent with the distribution of S-AUS across NSW (**Fig. 4**).

Interpreting MVEV circulation therefore requires consideration of both vector population structure and the broader ecological conditions in which transmission occurs. Whether the S-AUS and AUS lineages differ in traits that impact disease transmission, such as vector competence, seasonal abundance, meal choice or habitat use, remains unknown ^35^. Regional variation in MVEV vector competence in Australian *Cx. annulirostris* populations has previously been suggested, although mitochondrial lineages were not distinguished^44^. The 2023 outbreak followed two consecutive years of high rainfall linked to a La Niña weather event ^26,39^, conditions which may alter both mosquito and reservoir communities: increased availability of ephemeral water bodies provides breeding habitats for freshwater mosquitoes, including *Cx. annulirostris*, while also supporting waterbirds. Because *Cx. annulirostris* is an opportunistic feeder taking blood meals from both avian and mammalian hosts ^35^, it can act as a bridge vector for spillover infection into humans ^39^. Integrated surveillance of viral genomes, mosquito communities and vector population structure facilitates the investigation of these interacting processes.

As well as integrating pathogen and vector information, a further advantage of broad capture-based surveillance is the potential for detection of clinically important, but unexpected, viruses beyond the subset of pathogens that can be monitored with PCR-based testing. In this study, the recovery of SINDV led to its subsequent incorporation into ongoing PCR surveillance. LSV had never been detected in this region before and only nine complete genomes exist globally including the closely related Umbre virus. Two related isolates were recently implicated in two fatal encephalitis cases in France ^42^, and the virus has been shown to replicate in nerve cells. The co-detection of MVEV with SINDV and LSV further suggests that shared ecological conditions may support virus activity in these locations, which may provide a useful signal for future surveillance (**Fig. 4C)**.

Although COSMOS augments current surveillance methods, there are limitations of the approach and study design presented here. First, as in any mixed-catch approach, virus detections cannot be assigned to individual mosquito species within mixed pools, and targeted sampling or laboratory-based follow-up is likely required to clarify whether a particular species is contributing to transmission. However, COSMOS can be applied to individual mosquitoes; although this is necessarily less scalable than mixed-species surveillance, it can be particularly useful for characterisation of invasive or exotic mosquito specimens. Second, COX1-based relative abundance estimation may not directly reflect individual mosquito counts due to variation in mitochondrial copy number or other technical factors; nevertheless, estimates appear consistent with expected numbers for previously quantified species, and capture relative spatiotemporal changes in community composition (**Fig. 3**). Third, any compositional data, such as mosquito relative abundance, is sensitive to database composition and differential sensitivity for each component. Simulation-based benchmarking can help guide database design, but continued expansion and curation of mosquito reference genomes, and particularly sequencing of multiple individuals of genetically diverse species, is important for future applications. Fourth, the 219 samples analysed here were intentionally enriched for virus-positive traps, together with spatially and temporally matched negatives, to evaluate platform performance and arbovirus diversity. They therefore do not constitute an unbiased survey of mosquito or virus occurrence, and broader sampling across the surveillance program will be required to further resolve the patterns observed here. Finally, since targeted probe capture is inherently dependent on panel design, untargeted pathogens may be present in relatively low abundance in the metagenomic libraries after capture, and our bioinformatics workflow therefore included a check for the presence of additional viruses in background sequence (**Fig. S2**).

Overall, we find that pooled mosquito sampling can support substantially richer surveillance than is currently exploited, simultaneously resolving viral genomic diversity, mosquito community composition and cryptic population structure within vector species. Integrating these data layers allows emerging or circulating viruses to be interpreted within their ecological and population-genetic contexts, informing hypotheses about the drivers of outbreaks rather than documenting when they occur. As mosquito-borne disease patterns shift under changing climatic, ecological and anthropogenic pressures, scalable and integrated approaches such as COSMOS may be better positioned not only to detect emerging threats but also to understand the ecological contexts in which they arise and spread.

## Supplementary Materials

Methods, Supplementary Figures and Supplementary Results can be found in the enclosed ‘Methods and Supplmentary.pdf’. Supplementary Tables 1-6 are available in ‘Supplementary Tables 1-6.xlsx’.

## Data availability

The panel will be made available for purchase as a defined reagent through Twist Bioscience with design ID [TE_XXX]. Sequence data is available from the European Nucleotide Archive (ENA) with project ID PRJEB128300. Further data and analysis outputs are available for download from Zenodo (10.5281/zenodo.23310135).

## Code availability

A computational workflow designed to estimate mosquito species abundance within sequence data is available on GitHub (https://github.com/MultipathogenGenomics/COSMOS-abundance). All other associated analysis code is available in a separate repository on GitHub (https://github.com/MultipathogenGenomics/COSMOS-paper).

## Acknowledgments

We acknowledge all those involved in the NSW Arbovirus Surveillance and Mosquito Monitoring Program (ASMMP), which enabled this work.

