## Supplementary material for "COSMOS: integrated vector-pathogen metagenomics for mosquito-borne disease": Methods and Supplementary

### Supplemental

#### Methods

Study design and sample overview

This study describes the development and deployment of COSMOS, a custom targeted metagenomics sequencing approach designed to extend the NSW Health Arbovirus Surveillance and Mosquito Monitoring Program (hereafter, the NSW mosquito surveillance program). COSMOS was applied to archived cDNA extracts synthesised from 219 mosquito pools collected within traps, referred to here as surveillance samples. Virus detection and genome recovery, mosquito community composition, and mosquito population structure were assessed within surveillance samples as presented in the main text.

During COSMOS development, the approach was also applied to several control pools of mosquito specimens identified by an entomologist. In addition, a synthetic viral control and a serial dilution control of cultured Barmah Forest virus were used to assess viral target capture and quantitative recovery, respectively (**Supplementary Results**).

##### *Mosquito surveillance samples*

*Surveillance sample collection*

The NSW mosquito surveillance program captures and collects mosquito pools within carbon-dioxide baited light traps set regularly throughout the mosquito breeding season from November to April each year. Following collection, trap contents are maintained in cold chain during transport to the Department of Medical Entomology, ICPMR in Westmead hospital Sydney for further processing.

Individual mosquito numbers within each trap are counted, the trap contents homogenised, and total nucleic acids extracted from each mosquito pool and converted to cDNA according to in-house protocols. When resources permit, individual mosquitoes from a subset of traps are characterised morphologically prior to homogenisation to quantify distinct species [1].

*Routine PCR testing*

As part of the NSW mosquito surveillance program, detection of 8 pre-defined arbovirus species is carried out on all cDNA extracts. These arboviruses are: Edge Hill, Stratford, Japanese Encephalitis, Murray Valley Encephalitis, West Nile Kunjin variant, Barmah Forest, Kokobera and Ross River viruses.

*Surveillance sample selection*

In total, 219 cDNA extracts from surveillance samples underwent COSMOS enrichment. Surveillance samples were not selected randomly; 111 surveillance samples with positive PCR-based virus detections were selected, alongside a further set of 108 selected from a matched geographical location in the same year. Where an exact location-match was not available, a geographically nearby sample was selected. We aimed to sequence the majority of known arbovirus PCR-positives collected across each of the three seasons; a subset could not be located within freezer archives.

In total, we sequenced 42 traps from the 2021/22 season, 49 from the 2022/23 season and 128 from the 2023/24 season (**Fig. 1**).

***Mosquito pool controls***

*Single-species pools*

Sequence data derived from COSMOS enrichment of eleven mosquito pools was used to: (i) contribute to the reference database of mosquito COX1 sequences used in species assignment and (ii) act as sources for *in silico* mixtures of sequence reads used to validate computational mosquito species assignment from sequence data. These pools were composed of: *Aedes procax, Aedes multiplex, Culex sitiens,* members of the *Culex pipiens* complex, *Aedes vigilax, Culex annulirostris, Aedes notoscriptus, Coquillettidia linealis, Coquillettidia xanthogaster, Anopheles annulipes* and *Mansonia uniformis*. Several members of the *Culex pipiens* complex occur within Australia, many of which are challenging to distinguish morphologically. For simplicity, we refer to this as a "single-species" pool, although the *Cx. pipiens* pools may contain multiple species within the complex.

Mosquito specimens were sourced from field collections at Sydney Olympic Park. Mosquitoes were collected from carbon dioxide baited light traps and identified using taxonomic and pictorial guides. Whole mosquitoes were sorted into single species pools and nucleic acids extracted.

*Morphological identification pools*

A further pair of pools were also utilised to compare morphological identification with COSMOS-derived species assignment. Each pool was composed of ten individuals morphologically identified as *Culex australicus.* This mosquito is a member of the *Culex pipiens* complex and is morphologically very similar to other *Cx. pipiens* complex members but can be identified with significant expertise. These pools were not used in construction of the COX1 reference database.

*Laboratory mixture pools*

Two pools comprising four mosquito species were generated to test mosquito species identification and relative abundance estimation within mixtures. These were a mixture of *Aedes procax, Aedes multiplex, Culex sitiens* and *Culex pipiens* complex members (mixture 1) and a mixture of *Coquillettidia linealis, Aedes notoscriptus, Aedes vigilax* and *Culex annulirostris* (mixture 2). These were made by extracting DNA from 10 individual mosquito specimens separately per species using the identical extraction methods and mixing extracted DNA in equal volumes prior to library preparation. DNA mass was not normalised prior to mixing, in order to approximate routine workflow conditions.

*Laboratory serial dilution pools*

Ten serial dilution pools were generated using the DNA extracts described in the previous section, instead combined in unequal proportions. These pools were generated from two pairs of pool extracts: *Aedes vigilax* with *Culex annulirostris* and *Culex sitiens* with *Culex pipiens* complex members. The first pair was mixed in 1:99, 25:75, 50:50, 75:25 and 99:1 ratios and the second pair was mixed in 5:95, 25:75, 50:50, 75:25 and 95:5 ratios. Ratios refer to DNA extract volume contributed by each pair, and not normalised DNA mass.

***Virus sequencing controls***

*gBlock virus construct*

A synthetic construct of double stranded DNA representing 8 partial genome fragments from 7 virus species known as the gBlock control was sequenced to test COSMOS capture across viral targets. This was originally constructed as a positive control for PCR arbovirus detection performed by the NSW mosquito surveillance program. COSMOS capture of this sample was assessed in duplicate.

The construct is 646 nucleotides in length and contains a 66bp fragment of Ross River virus, a 73bp fragment of Barmah Forest virus, a 61bp fragment of Murray Valley encephalitis virus, a 70bp fragment of the West Nile virus Kunjin variant type 1, 107bp of the West Nile virus Kunjin variant type 2, an 82bp fragment of Kokobera virus, a 92bp fragment of Stratford virus and a 95bp fragment of Edge Hill virus.

*Barmah Forest virus culture*

A sample of Barmah Forest virus culture was sequenced to assess the quantitative relationship between virus abundance within samples (measured using qPCR cycle threshold) and the number of unique reads recovered by COSMOS.

An aliquot of culture was diluted in a four-fold series, and each dilution was tested for virus abundance using qPCR and underwent COSMOS enrichment.

Method details

***COSMOS capture probe design***

The COSMOS probe panel is a library of 48,043 biotinylated DNA probes of 120 nucleotides in length. These were designed to hybridise to:

- 116 arboviruses, defined here as viruses with a previously reported vertebrate and invertebrate host and prioritising those known to be mosquito-borne.
- 8 *Plasmodium falciparum* loci and their homologous regions within *P. vivax*, including cytochrome b, which is commonly used for *Plasmodium* species identification, and other loci associated with antimalarial drug resistance which have been targeted in previously published amplicon panels [2].
- 49 additional non-arboviral human viruses.
- 9 human pathogen bacterial loci.
- the mosquito cytochrome C oxidase subunit I gene (*COX1*), a mitochondrial gene that has been widely used as a species identification marker [3].

A complete list of targeted sequences is given in **Table S1**.

To design the COSMOS probe panel, sequences of desired targets were manually compiled using NCBI, including multiple individuals of each target species where available. The capture effectiveness of target enrichment has been shown to be consistently high when probe-target identity sharing is within at least 80% [4], allowing for the capture of within-species diversity. To ensure the COSMOS panel fell within this range, we clustered compiled sequences to 90% sequence identity using CD-HIT-EST [5] and generated 120bp probes using end-to-end tiling along a strict-majority consensus sequence. Low complexity sequences were then removed using self-self blast with dust filter enabled.

***RNA/DNA extraction and cDNA conversion***

For the 219 surveillance samples analysed in this study, remnant cDNA extracts stored by the NSW mosquito surveillance program were available. These were extracted using in-house protocols as previously described [1]. For single species pools and laboratory-based mixtures, total nucleic acids were extracted using the same method.

***qPCR detection of virus presence***

Virus-sample pair qPCR cycle threshold values for the surveillance samples were ascertained by the NSW mosquito surveillance program through routine testing at the time of their collection. The same method was used to test the Barmah Forest virus culture serial-dilution and g-block control samples shown in **Figure S3** and described in **Supplementary Results**. This method involves a series of multiplex fluorogenic TaqMan real-time RT-PCR assays with modifications, as previously described [1].

##### *Sequencing library preparation, hybridisation probe capture and sequencing*

All sequencing was performed within ICPMR in Westmead Hospital in Sydney. The workflow was based on the Twist Bioscience NGS target enrichment workflow with modifications. Briefly, cDNA extracts were used to prepare metagenomic sequencing libraries using the Twist EF1 library prep kit (Twist Bioscience) and the Twist Universal Adapter System, according to manufacturer directions. Libraries were pooled in equal volume and hybridised overnight with the COSMOS probe panel. Following COSMOS enrichment, captured fragments were amplified and cleaned using DNA purification beads.

The final library was then assessed for DNA concentration using the Qubit dsDNA High Sensitivity Quantification Assay and for fragment length distribution using the Agilent TapeStation, following manufacturer's protocols. Where multiple libraries were to be sequenced together, DNA mass and average fragment size were used to compute their loading proportions. Sequencing was performed on the Illumina MiSeq machine at 150bp paired end using 300 cycles, targeting approximately 1 million reads per sample.

#### Quantification and statistical analysis

##### *Sequence read processing*

Adapter sequences were removed from demultiplexed reads using Trimmomatic [6], and trimmed reads were used as inputs for genomic analysis software unless otherwise specified. Snakemake pipelines and Python scripts used for analysis can be found in an accompanying GitHub repository (https://github.com/MultipathogenGenomics/COSMOS-paper), as detailed in Code Availability.

##### *Virus analyses*

###### *COSMOS virus detection and genome recovery*

COSMOS virus detection and genome recovery was performed using the tool Castanet (v9.3) [7]. Castanet aligns reads across a user-defined set of input reference sequences that are assigned to target groups (hereafter the Castanet reference). Castanet collapses coverage to report a single read count and proportion of the target genome covered across each target group. This allows known diversity within a target group, e.g. multiple virus lineages, to be accounted for within the analysis. The Castanet reference matched sequences used to generate the COSMOS probe panel, which represented known diversity across the 116 targeted arboviruses.
A positive COSMOS virus detection was defined as ≥5% genome coverage at ≥2-fold read depth. Unique virus read count, that is the total number of virus reads present within a sample accounting for duplication during COSMOS enrichment, was also assessed. These were obtained from the 'prop_npos_cov2' and 'n_reads_dedup' columns in Castanet outputs respectively. These results are summarised in **Table S3**.

Virus sequence recovery was initially assessed as part of this study using a synthetic viral control and a serial dilution of cultured Barmah Forest virus, as described in **Supplementary Results** and shown in **Fig. S3.** The relationship between Ct and COSMOS-recovered unique virus read count across the four-fold serial dilution of Barmah Forest virus culture was assessed using ordinary least-squares linear regression implemented using linregress in SciPy (**Fig. S3**). Unique read counts were transformed as log10(read count + 1) prior to regression. Ninety-five per cent confidence intervals around the fitted regression line were calculated using Student's $t$ critical values with $n-2$ degrees of freedom.

##### *Virus consensus sequence generation*

Virus consensus sequences were generated by Castanet by implementing the –DoConsensus command line argument, which uses alignments across the target group within the Castanet reference. A minimum read depth of 2 was required to compute a consensus at a position, specified using the --ConsensusMinD argument. --DoKrakenPrefilter and –DoTrimming were also set to False alongside required Castanet command line arguments.

Little Sussex virus was treated separately because the Castanet reference did not contain a species-matched reference, although it did contain the related Umbre virus. For the single surveillance sample which was positive for Little Sussex virus, reads were realigned to concatenated segments of the Queensland Little Sussex virus sequence (NCBI Accession: OR077282-4) using minimap2 (v2.30-r1287) [8], converted to BAM file format using samtools (v1.22.1) [9] and a consensus made using ViralConsensus (v1.0.0) [10] using a minimum read depth of 2.

##### *Batch-specific handling of index hopping*

Sequencing was performed across four batches. One batch, which also contained the Barmah Forest virus dilution series, showed evidence of index hopping of BFV reads between libraries. For this batch the –PostFilt command line option was enabled. This removes reads with start-end coordinate pairs that are present only once within alignments, as these can arise through index hopping. For the remaining three batches, in which this pattern was not observed, --PostFilt was disabled because applying this filter reduced genome coverage for some virus-positive samples. To assess whether this batch-specific filtering influenced downstream results, virus detection and genome recovery within the affected batch were compared with --PostFilt enabled and disabled, as described in the **Supplementary Results** and shown in **Fig. S9**. The run with –PostFilt was performed using Castanet v9.2, unlike the rest of the runs which were performed with v9.3. Differences between these versions would not have impacted these analyses.

###### *Detection of insect specific viruses in surveillance samples from metagenomic background*

Hybridisation capture using COSMOS does not eliminate non-target sequence from libraries and therefore reads from insect-specific viruses infecting mosquitoes within surveillance samples were likely to be present within COSMOS sequence data. Using reads that were not aligned to COSMOS targets using Castanet, we performed assembly using SPAdes [11] with default parameters. Assembled contigs were assessed for similarity to the NCBI database using Kraken [12]. Virus assignments were filtered to give a set of additional viruses that were potentially present in the sequence data that were not targeted by COSMOS.

These viruses were manually compiled into a reference set containing 159 sequences representing 124 viruses. This reference set was combined with the COSMOS Castanet reference, and used as input to the tool Castanet, run as described above. For each virus in each surveillance sample, the number of unique reads and the proportion of each virus covered by ≥2 reads were assessed, as shown in **Figure S2** and described further in **Supplementary Results.**

The association between estimated *Culex australicus* relative abundance and Wuhan Mosquito Virus 6 unique read count across surveillance samples was assessed using ordinary least-squares linear regression implemented using linregress in SciPy (**Fig. S2**). Unique virus read counts were transformed as log10(read count + 1) prior to regression. Ninety-five per cent confidence intervals around the fitted regression line were calculated using Student's $t$ critical values with $n-2$ degrees of freedom.

###### *Relationship between PCR cycle threshold and virus genome recovery*

We assessed the relationship between viral abundance within samples and genome recovery using COSMOS in terms of both: (i) total recovered unique virus reads, and (ii) recovered genome coverage. PCR cycle threshold (Ct) values obtained from routine surveillance were matched to the corresponding proportion of genome recovered at ≥2-fold depth and the number of unique reads aligned to that virus genome, as reported by Castanet.

The relationship between virus qPCR cycle threshold (Ct) and COSMOS-recovered unique virus read count across surveillance samples was assessed using ordinary least-squares linear regression implemented using linregress in SciPy (**Fig. 3A**). Unique read counts were transformed as log10(read count + 1) prior to regression. Ninety-five per cent confidence intervals around the fitted regression line were calculated using Student's *t* critical values with $n-2$ degrees of freedom. For qPCR-positive sample–virus combinations for which COSMOS recovered no coverage, missing Castanet coverage values were interpreted as measured zero coverage.

The relationship between Ct and genome coverage was modelled using a four-parameter logistic function. Here, detections with positive Ct values but no virus coverage were included with a coverage value of 0. First, Ct values were grouped into 8 defined bins (0-16, 16-19, 19-22, 22-25, 25-28, 28-31, 31-34, 34-37), and median genome coverage values were computed within each bin. The model included lower and upper asymptotes, a midpoint and a slope parameter. The lower and upper asymptotes were constrained between 0 and 1, the midpoint was constrained to the observed median Ct range and the slope was constrained to be positive. Initial parameter values were set to 0, 1, the median Ct value across bins and at 1. Parameters were estimated from the binned median values using the curve_fit function within the SciPy package (v 1.17.1). Median genome coverage values within predefined bins were used for model fitting to reduce the influence of variable individual observations. From the fitted curve, we calculated the Ct at which predicted genome coverage was 50% and 95%, as displayed in **Figure 2.** Boxplot distributions of genome coverage within each bin are also shown **Figure S10**.

<u>*Phylogenetic analyses*</u>

Three virus phylogenies were generated as part of this work: Murray Valley encephalitis (MVEV), Little Sussex/Umbre virus (LSV) and a partial Sindbis (SINDV) phylogeny showing Genotype II, all shown in **Figure 4** in the main text. For each assessed virus, the virus name was searched for within the NCBI Virus database and all results downloaded in FASTA format. These sequences were combined with the COSMOS consensus output by Castanet or, in the case of LSV, generated using viralconsensus. These were aligned using MAFFT (v7.490)[13] using the –auto algorithm prior to tree estimation. Sequences were retained if at least 30% of their bases were not missing using the bash program awk.

For MVEV, 119 were available for phylogenetic analysis including nine sequences generated using COSMOS. Although ten MVEV detections were made in the surveillance samples, one detection had less than 30% virus coverage so was excluded.

Because only a single publicly available LSV genome was available, genomes of the closely related Umbre virus were included in the same analysis, producing a dataset of nine genomes. The three segments of the Little Sussex/Umbre viruses were manually concatenated prior to tree estimation.

For SINDV, the displayed phylogeny was restricted to genotype II, to which the COSMOS-derived sequences belonged.

All virus phylogenies were visualised using the R package ggtree [14].

##### *Maximum likelihood trees using IQTree*

For all three viruses, maximum likelihood phylogenetic trees were generated using IQTree [15] (v. 3.0.1) using the Generalised Time-Reversible (GTR) model (GTR+F+R6). These trees are displayed using the midpoint as the root, and tips ordered by increasing branch length.

##### *Time resolved trees and estimation of evolutionary rate using TreeTime*

For MVEV, TreeTime (v. 0.11.4) [16] was used to estimate a time resolved phylogeny from IQTree outputs using default settings and no specified prior on evolutionary rate. Results are shown in **Figure 4** and the root-to-tip regression output by TreeTime is shown in **Figure S11.** Dates were not available for 5/119 MVEV sequences, so these were estimated internally by TreeTime.

##### *Time resolved trees and estimation of evolutionary rate using BEAST*

For SINDV, Bayesian Evolutionary Analysis Utility (v10.5.0) was used to generate an xml file to construct a time-resolved phylogeny using Bayesian Evolutionary Analysis Sampling Trees (BEAST, v10.5.0). Using the output of this, and through secondary apps from the BEAST software package; Tracer was used to estimate the evolutionary rate and Tree Annotator was used to generate a Maximum Clade Credibility tree to evaluate sub-lineage emergence and temporal signal.

##### *Mosquito analyses*

<u>*Assessing enrichment of the COX1 gene*</u>

Data from the 219 surveillance samples was used to estimate enrichment of the COX1 gene described in the main text. First, all reads were aligned to a publicly available mitochondrial genome of the *Culex annulirostris* mosquito (NCBI accession: NC_054323.1), a common species in Australia. This was performed using minimap2 (v2.30-r1287) [8] and converted to BAM file format using samtools (v1.22.1) [9]. Next, per base read depth across the mitochondrial genome was computed using the bedtools (v2.31.1) [17] subcommand genomecov.

The position of the COX1 gene within this genome (1,430 - 2,966) was obtained from the NCBI genome annotation. To estimate an approximate enrichment ratio across *COX1*, we computed the median of the $\log_{10}$(read depth+1) across sites within and outside of the COX1 gene region for each sample, excluding a 300 base pair window flanking the COX1 gene. We then computed the mean of the sample-level medians for the COX1 and non-COX1 regions, and estimated enrichment as $10^{\bar{M}_{COX1}-\bar{M}_{non-COX1}}$, where $\bar{M}$ denotes the mean sample level median.

Because surveillance samples contained mixtures of mosquito species, reads were not expected to align uniformly to the *Culex annulirostris* mitochondrial reference. A single mitochondrial genome was used to provide a consistent coordinate system across all samples, with *Cx. annulirostris* selected because it was the most frequently detected mosquito species in the dataset. Divergence between non-Cx. annulirostris reads and the reference may have reduced mapping efficiency and led to an under-estimation of enrichment.

<u>*Estimating relative abundance of mosquito species in surveillance traps*</u>

Estimating mosquito relative abundance from COSMOS capture data followed two steps. First, each read was identified as belonging to a species based on assignment to a set of *COX1* reference sequences. Second, the relative count of reads assigned to each species was used to estimate relative abundance within the sample.

##### *Generating the mosquito COX1 reference set*

To compile a reference set, we collected *COX1* sequences from mosquito species found in the Indo-Pacific, concentrating on southeastern Australia [18]. Compilation of the COX1 reference database involved manual literature and NCBI searches, guided by Webb, Doggett & Russell

(2016) [18]. For some species, no sequence could be located, and in other cases only a partial COX1 sequence was available. We initially focused our compilation to sequences of at least 1,000 nucleotides in length (a typical mosquito COX1 gene is approximately 1,500 bp). Publicly available COX1 sequences were further supplemented using sequence data from the morphologically identified single-species pools. Because these were collected in Sydney, NSW, they were included to increase sensitivity to local mosquito populations.

We generated alignments of sequence data from these pools to reference COX1 mitochondrial genomes. Eight complete mitochondrial genomes were sourced from NCBI (*Aedes vigilax:* NC_027494.1*, Culex sitiens:* NC_054318.1*, Culex annulirostris:* NC_054323.1*, Aedes notoscriptus:* KM676219.1*, Coquillettidia linealis:* MN389455.*1, Anopheles annulipes:* NC_054312.1*, Mansonia uniformis:* NC_047479.1, *Culex pipiens molestus:* PQ584792.2) and reads from each pool were aligned to their matched reference using minimap2 (v2.30-r1287) [8] and converted to BAM file format using samtools (v1.22.1) [9]. For the three species for which a species matched mitochondrial genome was not available, a species of a matching genus was used (*Aedes procax* and *Aedes multiplex* reads were aligned to the *Aedes vigilax* reference*,* and *Coquillettidia xanthogaster* reads were aligned to *Coquillettidia linealis).* Alignment across the COX1 gene was generated using samtools (v1.22.1) [9], and the consensus sequence across this alignment using ViralConsensus (v1.0.0) [10] with a minimum read depth of 10.

Because available *COX1* references varied in sequence length and genomic coverage, sequences were restricted to a homologous region shared across the reference set. To achieve this, we generated a multiple sequence alignment using MAFFT (v7.526) [13], and trimmed nucleotides from either end where consecutive gaps occurred across the sequences. A small gap allowance of 3bp was permitted within each sequence to allow for INDELS within the sequence. Sequences that would reduce the homologous length to less than 500bp were also excluded. This produced a 931bp core region shared across available *COX1* sequences in the database.

Previous tests using simulations have shown that near-identical reference sequences can produce unexpected read assignments for the tool KMA [19]. We therefore removed sequences that were highly similar. This was performed by computing all 31bp kmers across the sequences and calculating pairwise mash distances. Mash distance is related to the Jaccard distance, which is the proportion of total kmers that are shared between a pair. Sequences were then clustered based on these distances using hierarchical clustering. Where mash distance < 0.01 between a collection of sequences, a representative sequence would be retained in the reference set. One representative sequence was retained from each cluster. Published reference sequences were prioritised over computationally generated consensus sequences. Among sequences with the highest available priority, the sequence with the lowest mean Mash distance to all cluster members was selected.

In general, this removed sequences from the same species. However, members of the *Culex pipiens* complex, which are considered different species (e.g. *Cx. quinquefasciatus* and *Culex pipiens*), were not sufficiently distinct across the 931bp fragment and were therefore clustered into a single analytical group. Here, we refer to this group collectively as the *Culex pipiens* group for simplicity. *Culex australicus,* which is also a member of the broader *Culex pipiens* species complex, was able to be distinguished using this COX1 fragment and remained a separate group.

Our final dataset consisted of 79 sequences representing 71 species spanning a homologous 931 bp fragment of the *COX1* gene. A list of sequences and their sources are given in **Table S4**. Sequence clustering used to construct the final database is shown in **Figure S12**, where those in bold were included in the final reference set.

It is worth noting that we observed that changing the reference sequences altered abundance estimates for some samples, highlighting the sensitivity of this type of analysis to representative sequence selection.

##### *Mosquito identification and abundance estimation using Kallisto and KMA*

To assign reads to COX1 reference set we used the tools KMA (v1.6.8) [20] and Kallisto (v0.51.0) [21]. First, we determined a species was present when both KMA and Kallisto exceeded a minimum assignment threshold of five reads or estimated counts, respectively. We further filtered results to references which had ≥90% coverage and at least 97.5% nucleotide identity across that region, as computed by KMA.

Where multiple sequences for a single species were present in the mosquito COX1 reference set, read counts were summed across the sequences, and the maximum template coverage and maximum nucleotide identity sharing across the covered region were used for filtering.

We estimated the relative abundance of identified species components using the numbers of reads assigned by KMA. For each sample, relative abundance was calculated as the number of KMA-assigned reads for each component species divided by the total number of assigned reads across all component species. The final species-identification and relative-abundance workflow was applied identically to the single-species pools, morphological comparison pools, laboratory mixtures, serial dilution pools and 219 surveillance samples.

Kallisto and KMA were run as follows:

<u>Kallisto</u>
First, an index was generated using the “index” subcommand and read assignment was performed using the “quant” subcommand using default command line arguments. Read counts on individual references were determined by the “est_counts” column in the resulting “abundance.tsv” file. For each assignment, 100 bootstraps were performed and results averaged across these iterations.

<u>KMA</u>
Index generation was performed using the “index” subcommand using default command line arguments. Read assignment was run in paired end mode (“-ipe”) using the additional command line arguments “-ef -ex_mode -vcf 2 -1t1”. The numbers of reads assigned to each reference was determined using the “readCount” value in the produced “.mapstat” file. For relative abundance estimation, KMA read counts were summed across references representing the same species and normalised by the total assigned count across retained species within each sample.

##### *Generating in silico mosquito mixtures for Kallisto and KMA benchmarking*

To evaluate whether COX1 read profiles could recover mosquito species composition, we generated *in silico* mixtures of reads produced from COSMOS capture of 11 single species pools.

To minimise estimation biases from other sources, we aimed to ensure that all reads within each mixture were reliably mosquito COX1 reads. First, reads in each single species pool were aligned to a set of three consensus sequences representing COX1 sequences across the genera *Aedes, Culex and Anopheles*, generated as part of the COSMOS probe panel design process. This was performed using minimap2 (v2.30-r1287) [8] and alignments converted to BAM file format using samtools (v1.22.1) [9]. Soft clipped bases were then removed from reads using code from a now deprecated repository called ngsutils (v0.5.9) [22] that we adapted for compatibility with Python3. Reads which fell entirely within the COX1 region were then extracted using bedtools (v2.31.2) [17] intersect and filtered to paired reads of at least 100bp in length. Reads were randomly sampled using the tool seqtk (v1.5-r133) [23] and combined in even proportions in every unique combination, producing 330 mixtures.

In order to test our computational workflow, reads in these mixtures were assigned to our database using the tools KMA and Kallisto (without using bootstraps). Results are shown in **Fig. S5** and described further in **Supplementary Results**. This includes a comparison of proportion estimation error between tools and the proportion assigned to an incorrect species.

##### *Validating relative abundance estimates in mixtures of morphologically similar taxa*

As described in the main text, we assessed relative abundance in two pools of ten individual mosquitoes which had been morphologically identified as *Cx. australicus*, but gave small proportions of abundance to the closely related *Cx. pipiens* complex group (**Fig. 3**). To assess whether this was due to uncertainty or because individuals were visually misidentified, we used *Cx. australicus* and *Cx. pipiens* complex COX1 references in the mosquito COX1 reference set (N=6) to determine species-defining polymorphic sites. These sites were defined as those where all references for that species carried the same allele which was entirely different to references for the other species.

Species-defining polymorphic sites were found using a multiple sequence alignment of the COX1 sequences made using MAFFT (v7.490)[13], ignoring sites where any reference carried a gap. Using minimap2 (v2.30-r1287)[8] and samtools (v1.22.1)[9], trimmed reads generated from each mixture were aligned to a single representative COX1 sequence (MN389460.1:1446-2982, which belongs to *Culex pipiens molestus*), and allele counts in read pile-ups were counted at each site using the tabulate-alignments tool within Iorek [24], using reads that were mapped with a minimum mapping quality score of 10. These counts were used to compute the fraction of reads at each site which carried the *Cx. pipiens* complex allele relative to all reads in the pile-up, as shown in **Figure S4.**

##### *Comparing estimates to surveillance program counts using concordance correlation coefficient*

We assessed whether our estimates of relative abundance were broadly in line with those published by the NSW mosquito surveillance program, which are based on morphological identification. Total mosquito counts, counts of *Aedes vigilax* and counts of *Culex annulirostris* are available for download from the NSW government website [25]. We matched the downloaded data to our samples using location and date, which matched 78 traps in total. 72 of these had reported *Ae. vigilax* counts and 75 had reported *Cx. annulirostris* counts.

Within the published count data, we computed the proportion of each species out of the total and compared this to our relative abundance estimates using Pearson's correlation and Lin's concordance correlation coefficient ($CCC$) [26]. $CCC$ assesses whether two sets of values sit on an $x = y$ line, i.e. are nearly identical. Using $x$ to represent proportions computed from morphological identification counts of either *Ae. vigilax* or *Cx. annulirostris,* and using $y$ to represent COSMOS-derived abundance estimates, we computed $CCC$ as follows:

$$CCC = \frac{2cov(x,y)}{\sigma_x^2 + \sigma_y^2 + (\bar{x} - \bar{y})^2}$$

Where $cov(x,y)$ represents the covariance between $x$ and $y$, $\sigma_x^2$ and $\sigma_y^2$ represent the variances and $\bar{x}$ and $\bar{y}$ represent the means. In this formula, the numerator is therefore large when $x$ and $y$ vary together, while the denominator penalises differences between the two sets. Values approaching 1 indicate high concordance.

<u>*Assessment of mosquito community composition*</u>

##### *The principal components of mosquito community composition*

A principal component analysis (PCA) was performed on 213 samples which carried mosquito abundance estimates. In this analysis, we split the estimated abundance of *Cx. annulirostris* into S-AUS and AUS lineages, which is described later within **Methods**. We treated these lineages separately, thus computing PCs over 24 groups. On **Figure 4** in the main text, and **Figure S6**, we consider lineage proportions in samples where all four lineage informative sites were covered with at least 20 reads. Of 170 *Cx. annulirostris* detections, 157 met this criterion. Rather than excluding the remaining 13 samples from the PCA, the proportion estimate was used in these samples such that they could be retained. Although estimates from these lower-coverage samples are expected to be less precise, allele proportions across the four lineage-informative sites were generally concordant. To assess whether inclusion of these samples influenced the PCA, the analysis was repeated after excluding the 13 samples that did not meet the high-confidence coverage threshold. PC scores were nearly identical between the two analyses, linear regression of scores from the sensitivity analysis against those from the primary analysis gave slopes of 1.000 for PC1 ($R^2$ = 0.999) and 0.999 for PC2 ($R^2$ = 0.996), and loadings were similarly correlated between each analysis ($R^2$=0.999 and 0.997 for PC1 and PC2 respectively), indicating that inclusion of the lower-coverage samples had little influence on the overall community analysis.

To account for the compositional nature of this data type, we first used the Hellinger transform, which standardises abundances by total for each sample (which has no effect here as all sum to 1) and square roots the values. We performed this using the Python package NumPy. We then performed PCA using 5 components using the package scikit-learn.

A summary figure showing scatter plots of each sequential PC, variance explained and loadings is shown in **Figure S13**.

##### *Environmental and temporal correlates of community composition*

To assess the correlates of environmental and temporal factors with community composition, we fitted separate linear mixed-effect regression models using the Python package statsmodels. Each model included a predictor variable and sampling site as a random intercept to account for repeated sampling at the same location. PC1 and PC2 values were included as outcomes. Each regression followed the general formula:

$$PC \sim variable + (1|\, location - site)$$

Across the 213 tested traps there were 108 unique sites.

Categorical variables were encoded against a reference category as detailed below. Continuous predictors were standardised by subtracting their mean and dividing by standard deviation so that estimated effect sizes represent the change in outcome associated with one standard deviation change in the predictor. Distance to coast was log10-transformed (including a small pseudocount of $10^{-10}$) prior to standardisation.

The tested variables were as follows:

| ***Predictor*** | ***Transformation or coding*** | ***Categories*** | ***Reference category*** |
|---|---|---|---|
| ***Environmental*** | | | |
| *Location type* | Categorical | Metropolitan fringe, Metropolitan, Rural, Urban | Metropolitan fringe |
| *Position relative to the Great Dividing Range (GDR)* | Categorical | East, West | East |
| *Distance to coast (km)* | $\text{Log}_{10}$, followed by z-score | — | — |

| | | | |
|---|---|---|---|
| *Elevation (m)* | z-score | — | — |
| *Rainfall in the three weeks before collection (mm)* | z-score | — | — |
| ***Temporal*** | | | |
| *Sampling season* | Categorical | 2021–22, 2022–23, 2023–24 | 2021–22 |
| *Sampling week* | z-score | — | — |

Across categorical variables, overall associations were tested using the Wald chi-square test, which are reported on **Figure 5** in the main text.

Prior to testing, PC1 scores were multiplied by -1 so that it roughly was oriented in the west to east direction. Loadings were retained using their original sign.

To correct for multiple testing, we used the Benjamini-Hochberg correction implemented in statsmodels and report these p-values. This was performed across 22 comparisons which included the tests listed in the table above as well as models testing the variables: number of mosquito species detected within each trap, whether a virus was detected in the trap, Simpson's diversity index ($D_1$) and Office of Local Government council designation code; these results were not depicted in the main text, but are reported in **Table S5**.

<u>Environmental and spatial variables</u>

Coastline information was obtained from the Geoscience Australia website as a shapefile [27]. Latitude and longitude of each trap was determined using the given location in received metadata. Distance from coast was calculated as the shortest distance between each trap location and the coastline layer using the Python package geopandas (v1.1.3). The coordinates of the GDR were found using Google Earth, where a reference map was used to trace the GDR line and extracted as a JSON file. These coordinates are also used in NSW map visualisations and to estimate whether a surveillance sample fell east or west of this divide.

Whether a trap was collected in a rural/urban/metropolitan-fringe or metropolitan region was determined from the NSW government website [28], based on the council area within which the longitude and latitude for the trap fell.

Trap elevation and the rainfall in the preceding three weeks to trap collection were extracted from the Open-Meteo Archive API using the openmeteo_requests Python package. For each trap, total precipitation was calculated over the 21-day period ending on the sampling date by summing daily precipitation values (precipitation_sum) returned for the corresponding latitude and longitude coordinates.

<u>Temporal variables</u>

Collection season and week were available as surveillance sampling metadata.

##### *Comparing diversity between surveillance samples using Simpson's diversity index*

To compare species diversity across surveillance samples we used Simpson's diversity index ($D_1$) [29]. For each surveillance sample, let $p_i$ represent the relative abundance of species $i$. The species diversity is therefore:

$$1 - D = 1 - \sum p_i^2$$

This is the probability that two randomly selected individuals from this trap are different species, under the estimated relative abundance distribution. This spans from 0 to 1 where values closer to 1 indicate greater diversity.

Because multiple surveillance samples were collected from the same site within the same season in our dataset, we performed a linear mixed-effects model comparing $D_1$ with distance to coast ($\log_{10}$ transformed) across metropolitan, metropolitan-fringe, urban and rural regions, using sample site as a random intercept. The results are shown in **Figure S7**.

*Assessment of cryptic population diversity in Culex annulirostris*

Mitochondrial lineages of *Cx. annulirostris* have been previously detected using partial sequences of the *COX1* gene [30]. 208 publicly available sequences of a 538bp COX1 fragment have been annotated as one of five distinct lineages: Australian (AUS; N=122), South Australian (S-AUS;N=15), Papua New Guinea 1 (PNG-1; N=37), PNG-2 (N=19) and Solomon Islands (SI; N=15) [30].

COSMOS also enabled assessment of cryptic population structure in additional mosquito species. As an example, we show an analysis of *Aedes notoscriptus* based on previously published sequences [31] (**Supplementary Results**). This was performed using the same method as the *Culex annulirostris* analysis, except that we designated lineages based on a threshold of 97% identity sharing, which generated three lineages here named population 1-3.

##### *Using lineage informative kmers to infer presence of multiple Culex annulirostris lineages in surveillance traps*

To assess for evidence of the non-Australian *Cx. annulirostris* PNG-1, PNG-2 and SI lineages in surveillance samples we computed 31 bp kmers across the published sequences and compiled the sets of kmers which appeared in at least 95% of individuals within each lineage, but in no sequence belonging to another lineage.

We used JellyFish (v1.1.12) [32] to then count all 31bp kmers within trimmed sequence reads across our surveillance traps. Counts for the lineage-informative kmers were extracted, summed and plotted in **Figure S14**. We did observe small counts of certain kmers within traps: five samples had 1-2 counts of one to two SI specific-kmers, one sample had 1 count of two PNG-1 specific kmers and there were no counts of PNG-2 kmers in any trap. Kmer matching is conservative and may fail to detect divergent individuals within a lineage. However, low counts across a set is more likely to represent sequencing error or incomplete lineage sorting, and we therefore did not interpret this as evidence of lineage presence.

##### *Identifying mitochondrial lineage proportions of Culex annulirostris using lineage defining sites*

To estimate the relative proportions of the *Cx. annulirostris* S-AUS and AUS lineages in the surveillance samples, we identified lineage-defining single nucleotide polymorphisms between these groups in the publicly available sequences. Because we observed minimal evidence of the PNG-1, PNG-2 and SI *Cx. annulirostris* lineages using kmers, these lineages were excluded from this analysis.

A site was identified lineage-defining when a single nucleotide was entirely shared within the AUS lineage while being entirely absent from the S-AUS lineage. To identify these sites, we generated multiple sequence alignments of the published partial sequences and scanned these for sites which satisfied this definition. Sites containing any gaps were excluded.

This identified 4 lineage-informative sites for *Cx. annulirostris* AUS/S-AUS. In reference sequence co-ordinates, these were: 1,960 (S-AUS: T; AUS: C), 1,987 (A; T), 2,017 (T; A/G), and 2,077 (T; A) within the *Cx. annulirostris* mitochondrial reference genome (NC_054323.1).

We aimed to assess only read pairs that likely belonged to *Cx. annulirostris* within the surveillance samples. We identified these reads based on read assignment to any *Cx. annulirostris* reference within the mosquito COX1 reference set, which overlaps the 538bp fragment used to identify lineage defining sites. Read identifiers were obtained from the ".frag.gz" file produced by KMA and reads were extracted from trimmed sequence data files using seqtk (v1.5-r133) [23] subseq. These reads were then re-aligned to a whole genome mitochondrial reference sequence (NC_054323.1) using minimap2 (v2.30-r1287) [8] and converted to BAM file format using samtools (v1.22.1) [9].

The position of lineage-defining sites within fragments was converted to reference sequence coordinates by finding the location of the first and last 31 bp kmer in a representative fragment ((DQ673677.1 ) using BLAST (2.16.0+) [33].

To estimate relative proportions of each we used read pile-ups. For each sample, base calls within reads covering each lineage-defining position with a mapping quality of at least 20 were counted using the tabulate-alignments tool within Iorek [24].

For each site, the proportion of reads carrying the S-AUS lineage-defining alleles were calculated relative to total pile-up count at that position, and final lineage proportions estimated by averaging these site-level proportions across informative sites. Proportion estimates were rounded to three decimal places prior to use in subsequent analyses.

To assess whether lineage-defining alleles co-occurred on the same read pairs, linkage between consecutive sites was evaluated using the Python package Pysam (v0.23.3). For each consecutive pair of sites, read pairs covering the first position and the second position were assessed for base identity at each position. Counts of each combination of alleles for each pair (i.e. 2x2 haplotype count table) were used to compute Pearson's correlation between sites. We restricted this to pairs of sites within a sample with at least 20 read pairs in the pile-up. To limit computation, each site-pair comparison within a sample was capped at 500 alignment records reaching the end of the filtering procedure. Each read pair contributed to at most one site-pair comparison within that sample.

For inclusion in **Figure 4** in the main text, all four *Cx. annulirostris* sites were required to have a pile-up depth of ≥20 reads with a mapping quality of ≥20 (N=157/170). Because many sampling sites overlap, on **Figure 4** we opted to average S-AUS proportion across time points. Latitude and longitude values were rounded to two decimal places (roughly within 1 kilometer) and grouped based on these values. 157 observations were thus grouped into 78 points on the plot. Grouping information and computed results can be found in **Table S6.**

The relationship between sample distance from the coast and the estimated proportion of the S-AUS *Cx. annulirostris* lineage was assessed using ordinary least-squares linear regression implemented using linregress in SciPy (**Fig. S6**). Linear regression was used as an exploratory analysis, recognising that the bounded nature of lineage-proportion data may not fully satisfy the assumptions of linear regression. Ninety-five per cent confidence intervals around the fitted regression line were calculated using Student's $t$ critical values with $n-2$ degrees of freedom.

The relationships between latitude and the estimated relative proportions of *Ae. notoscriptus* population 1 versus populations 2 or 3, and population 2 versus population 3, were assessed using ordinary least-squares linear regression implemented using linregress in SciPy (**Fig. S8**). Linear regression was used as an exploratory analysis, recognising that the bounded nature of population-proportion data may not fully satisfy the assumptions of linear regression. Ninety-five per cent confidence intervals around the fitted regression lines were calculated using Student's $t$ critical values with $n-2$ degrees of freedom.

##### *Spatial analysis and visualisation*

A NSW map file was downloaded from the Australian Bureau of Statistics website in GeoPackage format [34]. Maps and all other plots were generated using matplotlib (v3.10.7) and seaborn (v0.13.2) in Python.

Within **Figure 4** in the main text, coastal and non-coastal was determined based on whether the trap was >20km from the NSW coastline. This threshold is approximately equivalent to that used by the NSW mosquito surveillance program to designate traps.

**Supplementary Figures**

Figure S1

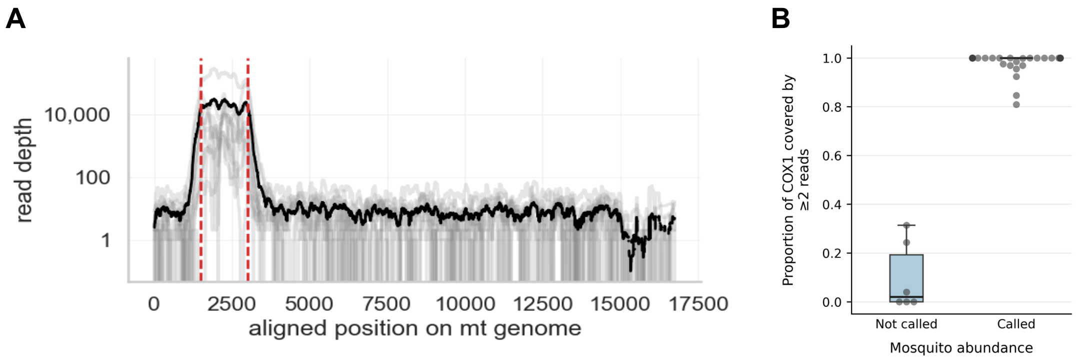


**Figure S1. Performance of COSMOS mosquito COX1 capture across single-species control pools and surveillance samples. (A)** Each line represents a single mosquito species pool that was enriched using COSMOS. The x-axis shows a position within a multiple sequence alignment of mosquito mitochondrial genomes, and the y-axis shows the read-depth of sequence reads aligned to each position. Reads from each pool were aligned to a reference sequence of a matching species, or a species in the same genus where a reference was unavailable, as detailed in **Methods**. The black line shows the median read depth across pools, and the red dashed lines indicate the targeted *COX1* gene region. **(B)** Proportion of the COX1 targets covered at ≥2-fold depth (y-axis) for surveillance samples with and without mosquito species relative abundance calls from the COSMOS computational workflow (x-axis). Of the 219 surveillance samples, six do not have species calls (‘Not called’). These samples also showed low *COX1* recovery. Coverage was calculated using Castanet against the mosquito consensus reference sequences described in **Methods**.

Figure S2

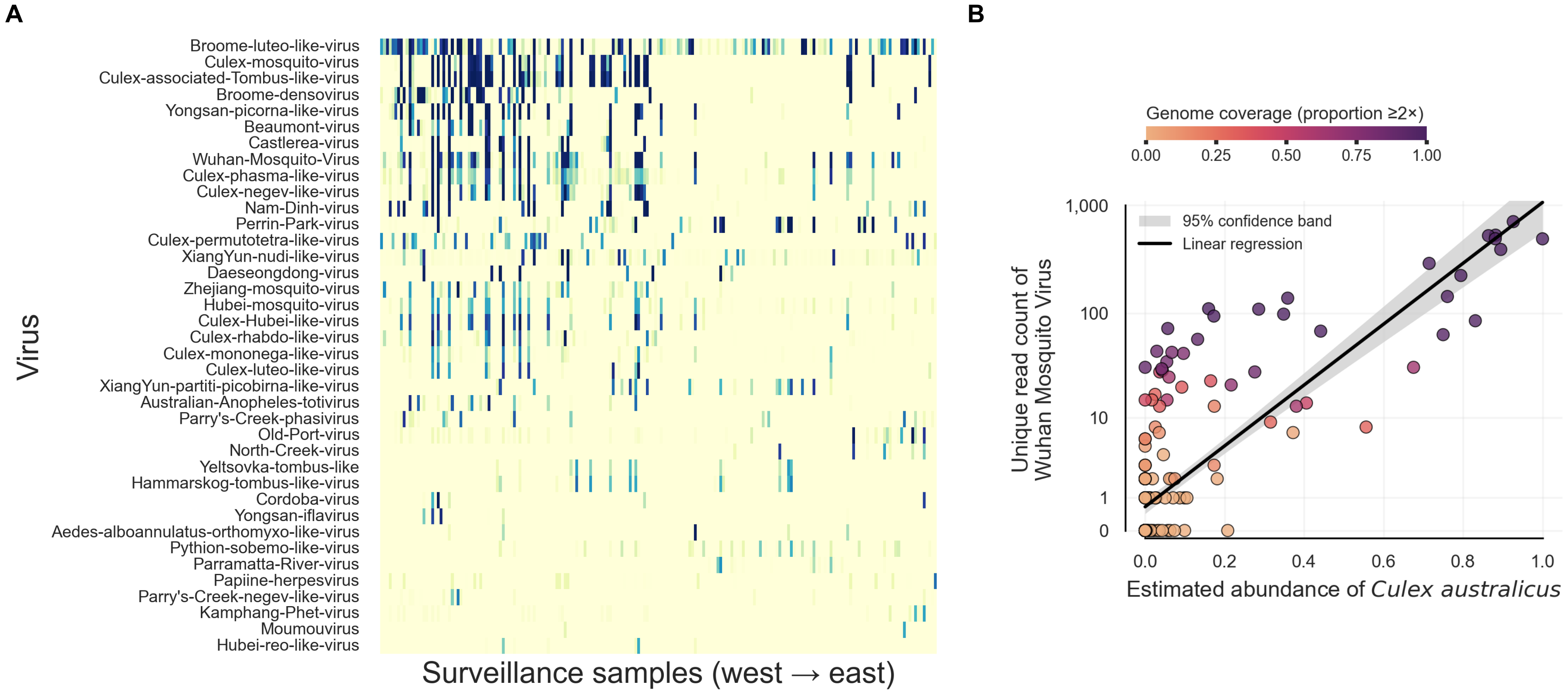


**Figure S2. Mosquito virome recovery from metagenomic background in surveillance samples. A)** Insect specific viruses detected across surveillance samples. Rows represent viruses with at least 50% genome coverage in at least one sample (N=38), and cell colour indicates the proportion of the viral genome covered at ≥2-fold depth. Surveillance samples are ordered west to east. **B)** Association between estimated *Culex australicus* relative abundance (x-axis) and Wuhan Mosquito virus unique read count (y-axis; $\log_{10}$(read count + 1)) across surveillance samples with mosquito abundance estimates (points; N=213). Points are coloured by proportion of the virus genome covered by at least 2 reads. A linear regression was fitted with the model $\log_{10}$(read count + 1) = 0.50 + 6.46 × *Cx. australicus* relative abundance ($R^2$=0.63, P=5.22 $\times 10^{-47}$, N=213).

Figure S3

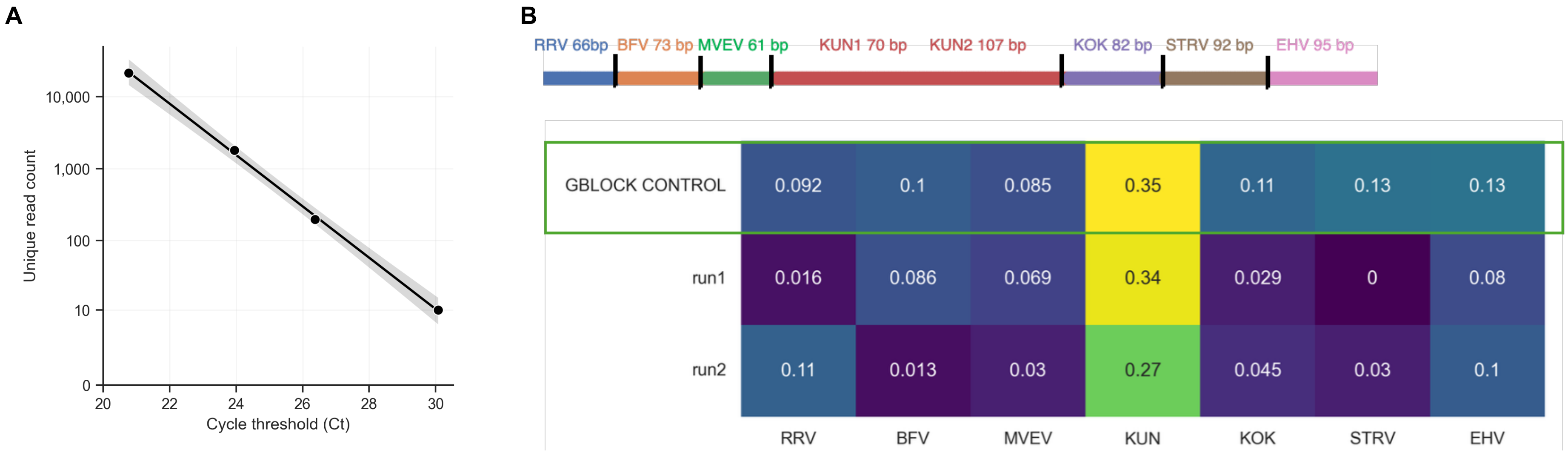


**Figure S3. COSMOS-recovery of virus controls.** (A) Points show a four-fold serial dilution of Barmah-Forest virus culture that were enriched using COSMOS. The x-axis shows the cycle threshold determined via qPCR of the diluted sample and the y-axis shows the number of unique virus reads counted within produced sequence data using the tool Castanet. A linear regression was fitted with the model: $\log_{10}$(unique read count + 1) = 11.83 - 0.36 $\times$ Ct ($R^2$=0.999, N=4, P=0.0004). (B) The top shows a schematic of the synthetic construct annotated with virus names (RRV: Ross River virus, BFV: Barmah Forest virus, MVEV: Murray Valley Encephalitis virus, KUN1: Kunjin virus variant 1, KUN2: Kunjin virus variant 2, KOK: Kokobera virus, STRV: Stratford virus, EHV: Edge Hill virus) and partial genome lengths. The top row of the heatmap shows relative lengths of each virus (columns) within the construct, as ordered within the schematic. The two following rows show the relative number of reads recovered for each virus across two separate sequencing runs following COSMOS enrichment.

#### Figure S4

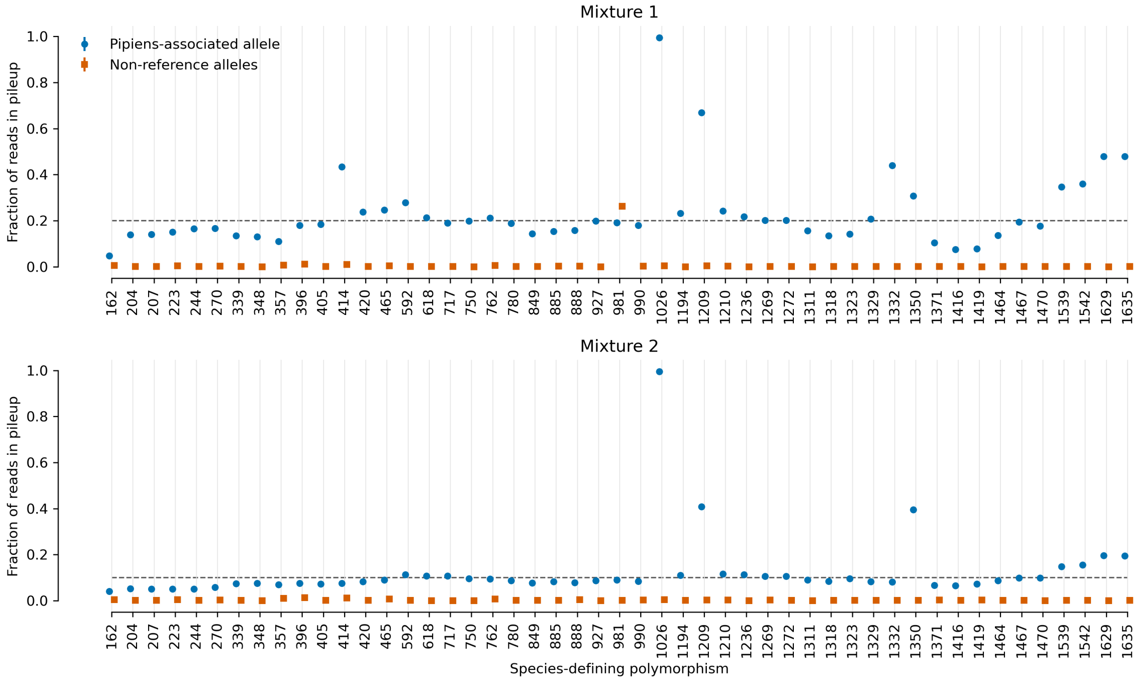


**Figure S4. Allele proportions at COX1 polymorphisms distinguishing *Culex australicus* from other *Culex pipiens* complex members in morphologically identified mosquito pools.** Read pile-ups were assessed at COX1 sites identified as polymorphic between *Cx. australicus* and other members of the *Cx. pipiens* complex determined using references included in the mosquito relative abundance estimation database. At each site, points show the proportion of reads carrying the *Culex pipiens* complex allele (blue) and other alleles (orange) relative to the total pile-up count. Each panel shows a mixture of ten individual mosquitoes that were morphologically identified as *Cx. australicus.* Across sites, the proportion of *Cx. pipiens* complex alleles were consistently approximately 0.2 (mixture 1) and 0.1 (mixture 2), consistent with the relative abundance estimates obtained from the COSMOS workflow. A small number of sites deviate, which may reflect that these sites were not species-defining which may be expected given the limited number of references used to define these sites (see **Methods**).

Figure S5

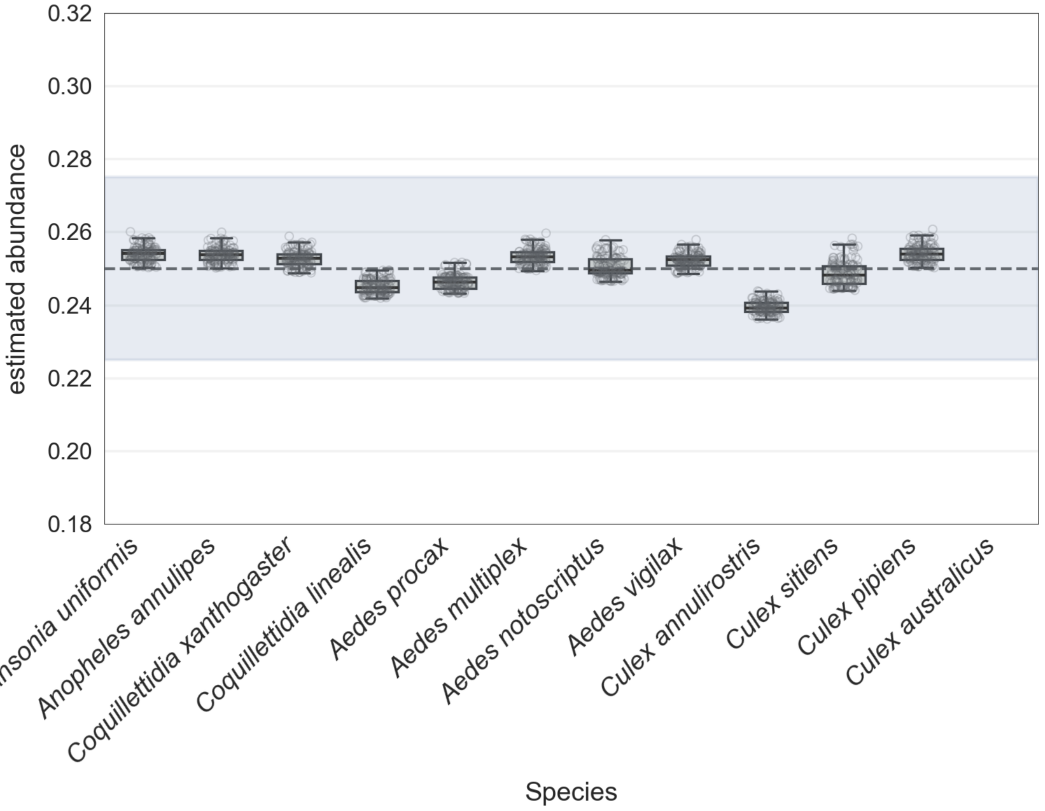


**Figure S5. Mosquito relative abundance estimation across *in-silico* mixtures of reads.** Every four-species combination of 11 single species pools were generated in equal proportions (N=330). Prior to mixing, reads were aligned to three consensus sequences of COX1, soft clipped bases were removed, and reads were filtered to a minimum length of 100bp, as described in Methods. Proportions of each species were then estimated using our workflow. Boxplots show the distributions of estimated proportions (y-axis) across the species included in the mixtures (x-axis). The blue shaded region shows a ±10% relative error around the expected abundance of 25%.

Figure S6

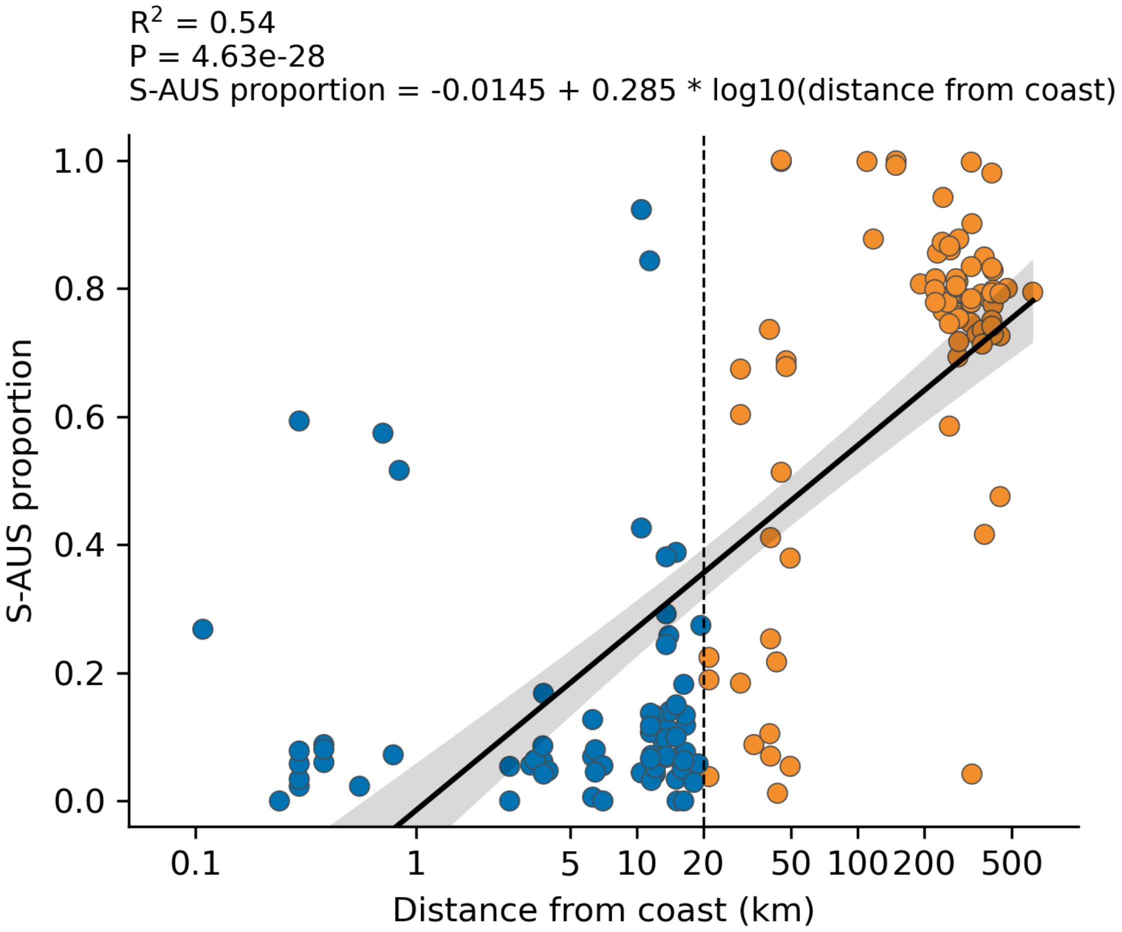


**Figure S6. Relative proportion of the *Culex annulirostris* COX1 mitochondrial lineage S-AUS increases with distance to coast.** Each point represents a surveillance trap with a positive detection of *Cx. annulirostris*. S-AUS proportions were estimated from read pile-ups at lineage-defining polymorphic sites. Distance from the coast is shown in kilometres on a $\log_{10}$ scale. Points are coloured as coastal (<20 km) or inland (≥20 km); the dashed vertical line marks 20 km which is typically used to denote coastal and non-coastal in NSW. The solid line shows an exploratory linear regression, and shading indicates the 95% confidence interval for the fitted mean: S-AUS proportion = −0.01 + 0.285 × $\log_{10}$(distance from the coast in km) ($R^2 = 0.54$, $P = 4.63\times 10^{-28}$; $n = 157$). The linear fit describes the association but does not constrain predicted proportions to 0–1.

Figure S7

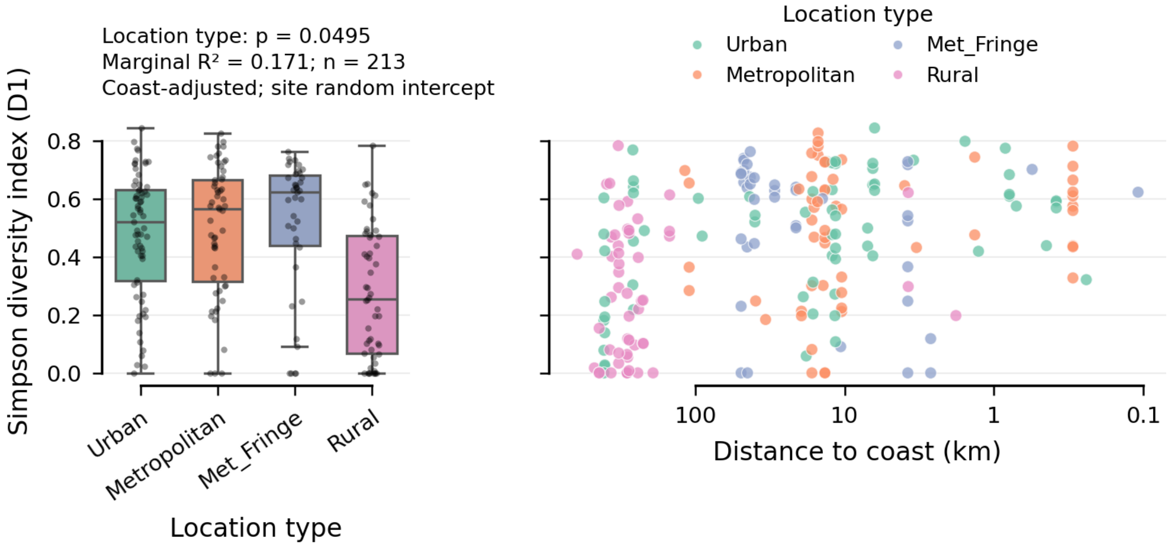


**Figure S7. Mosquito species diversity between location types.** Boxplots on the left show Simpson's diversity ($D_1$) across surveillance samples between location types, which were grouped according to government assigned codes for each council. The scatter on the right shows distance to coast (on a log scale; x-axis) against diversity ($D_1$; y-axis). Text annotations describe a mixed-model regression comparing $D_1$ across the four group categories, including sampling location as a random effect to account for repeated sampling.

Figure S8

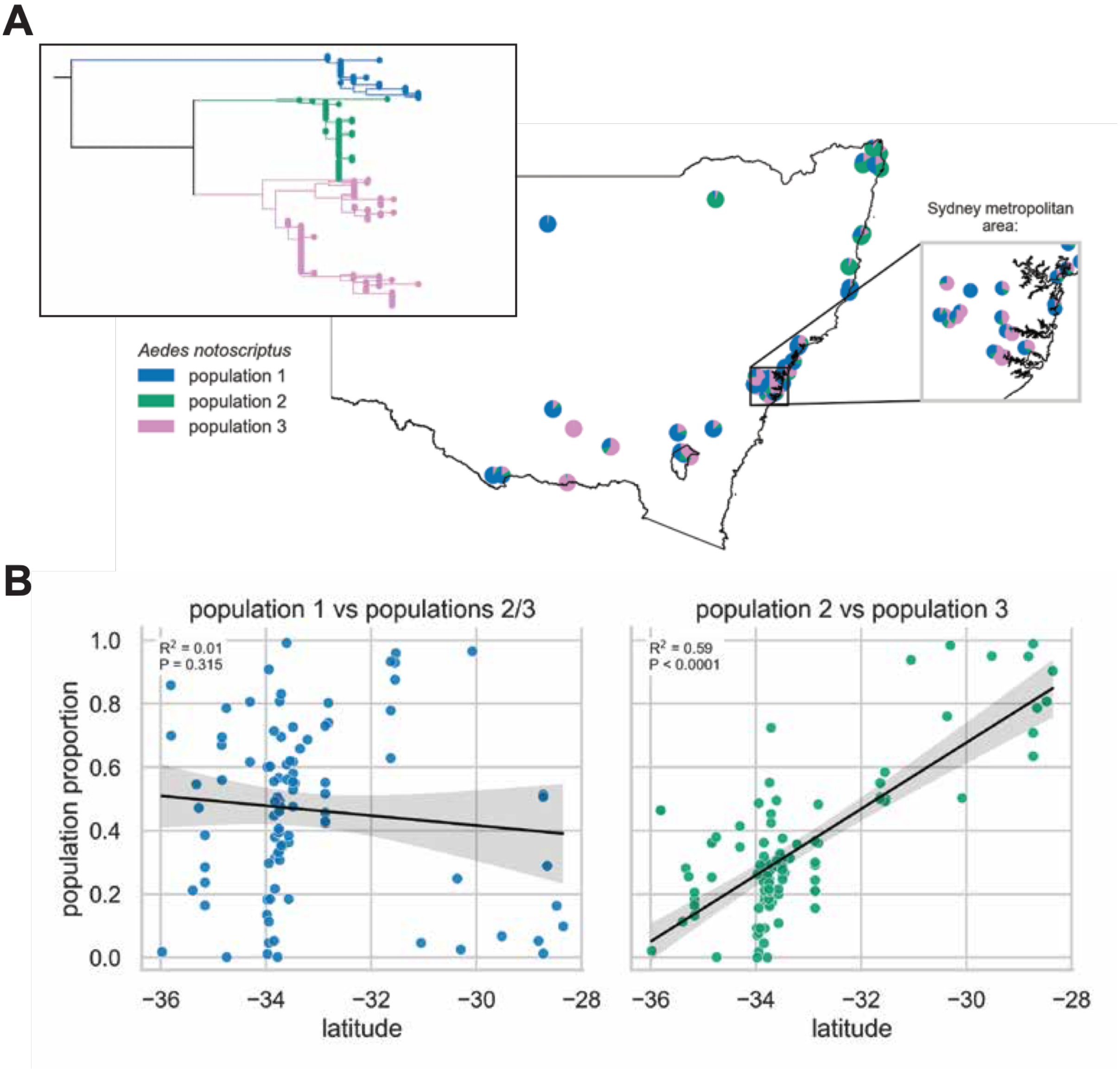


**Figure S8. The distribution of three cryptic mitochondrial *COX1* subpopulations of *Aedes notoscriptus*. A)** Each pie chart represents a surveillance trap and shows the location and relative proportions of three *COX1* subpopulations of *Aedes notoscriptus*, as indicated by the figure legend. Proportions are reflected as pie segments, which were computed using read pile-ups at population defining sites identified from publicly available sequences published by Endersby *et al.* (2013). The Inset shows the phylogenetic trees for these sequences, coloured to match their assigned population. Populations were defined from a pairwise distance tree using the publicly available partial *COX1* sequences [31]. Traps were retained if they had at least 12/15 sites which differentiated population 1 from 2 or 3, and 5/6 sites which differentiated 2 and 3 for *Ae. notoscriptus* (N=97). **B)** Points show a surveillance trap with a positive identification of *Ae. notoscriptus.* The x-axis shows the latitude (position on the north-to-south axis) and the y-axis shows the relative proportion of population 1 vs. 2 or 3 (left), and population 2 vs 3 (right). Read pile-ups at population defining polymorphic sites were used to compute relative proportions of each population, with a set of sites used to differentiate population 1 from the more closely related 2 and 3 groups, then a second set used to differentiate population 2 and 3. For each, a linear regression was performed which fitted the models population 1 vs 2/3 proportion = - 0.05 - 0.02 $\times$ latitude ($R^2$=0.01, N=97, P=0.32) and population 2 vs 3 proportion = 3.8 + 0.10 $\times$ latitude ($R^2$=0.59, N=97, P=4.07 $\times 10^{-20}$).

Figure S9

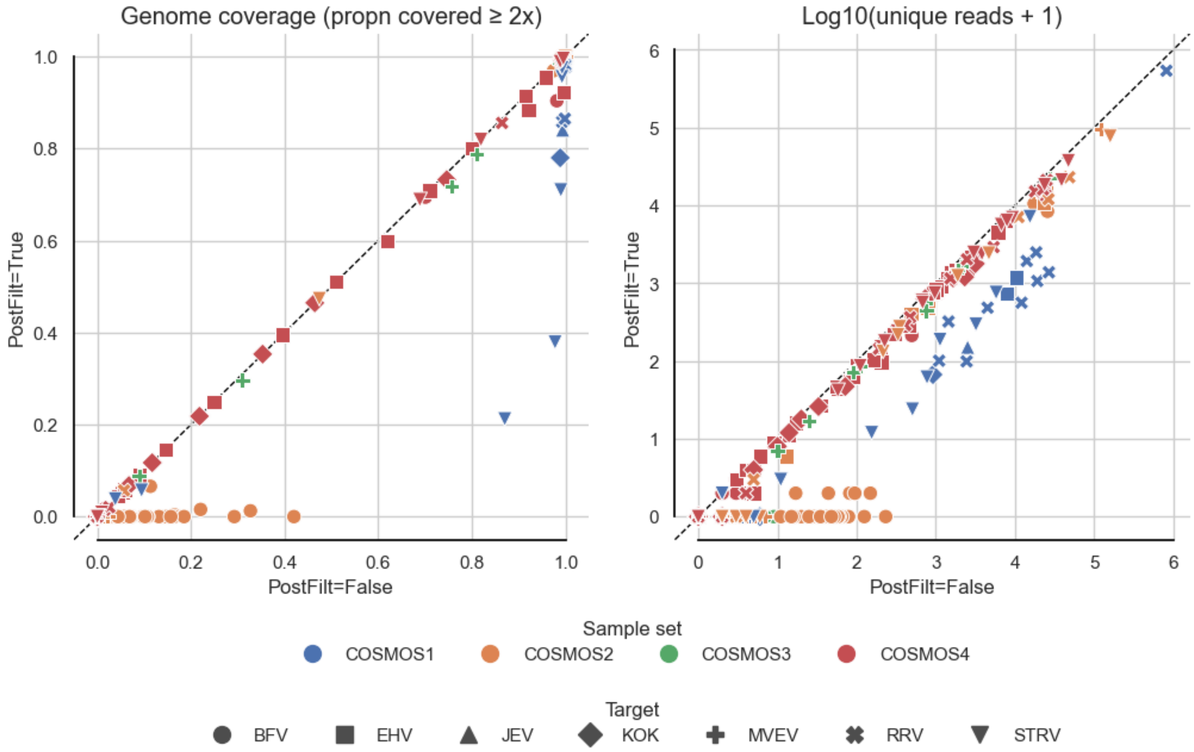


**Figure S9. Impact of –PostFilt on virus genome recovery reported by Castanet.** Virus genome coverage at ≥2-fold depth (left) and unique virus read count (right) reported by Castanet with (x-axis) and without (y-axis) the use of the command line argument –PostFilt. Sequencing of 219 surveillance samples was performed over four batches named COSMOS1-4, here indicated by marker colour. Marker shape indicates virus identity. In COSMOS2, low level BFV signal was observed only when –PostFilt was not applied, consistent with index-hopping from BFV-positive libraries within this batch. In contrast, COSMOS1 showed reduced genome coverage for otherwise high coverage detections when –PostFilt was applied. Results are described further in **Supplementary Results.**

Virus acronyms: EHV, Edge Hill virus; STRV, Stratford virus; RRV, Ross River virus; KOK, Kokobera virus; MVEV, Murray Valley encephalitis virus; BFV, Barmah Forest virus; JEV, Japanese encephalitis virus.

Figure S10

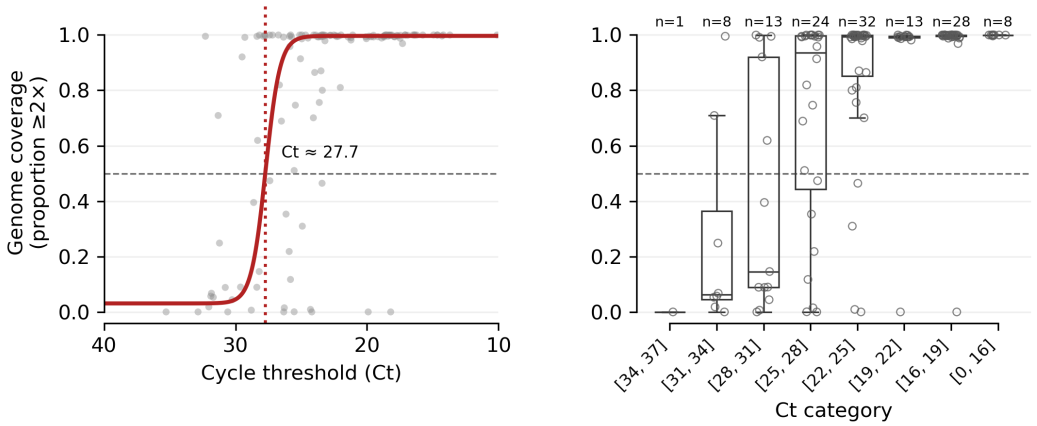


**Figure S10. The relationship between COSMOS-recovered virus genome coverage and PCR cycle threshold value.** PCR cycle threshold (Ct) values from routine surveillance were grouped into predefined bins and the median COSMOS-recovered virus genome coverage at ≥2-fold depth was computed within each bin. Virus-sample pairs with a positive Ct value but no recovered virus genome were assigned a genome coverage value of 0. (A) A four-parameter logistic model of the relationship between Ct value and recovered virus genome coverage. Parameters were estimated using median Ct and median genome coverage within each bin as described in Methods. The vertical red dotted line indicates the Ct at which the fitted curve predicted 50% genome coverage (27.7). (B) Box plot distributions of genome coverage within each bin. Number of detections in each bin are shown above each box. In A and B, Ct axes have been reversed such that higher viral abundance is shown from left to right.

Figure S11

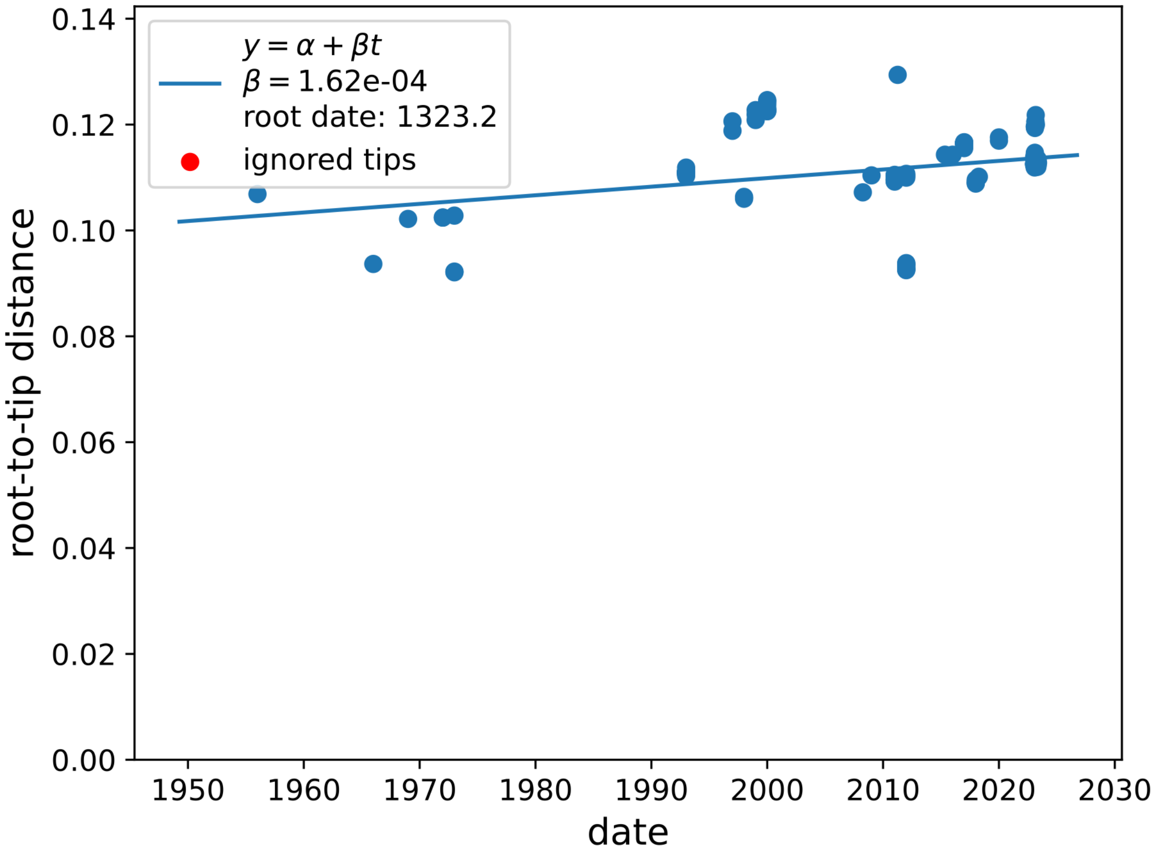


**Figure S11. Root-to-tip regression from a molecular clock and time-resolved phylogeny analysis of MVEV using Treetime.** Each point represents a sequence, downloaded directly from Genbank on NCBI Virus or generated using COSMOS (N = 119). The mutation rate observed from these sequences was 1.624e-04 and the coefficient of determination (r2) was found to be 0.09.

Figure S12

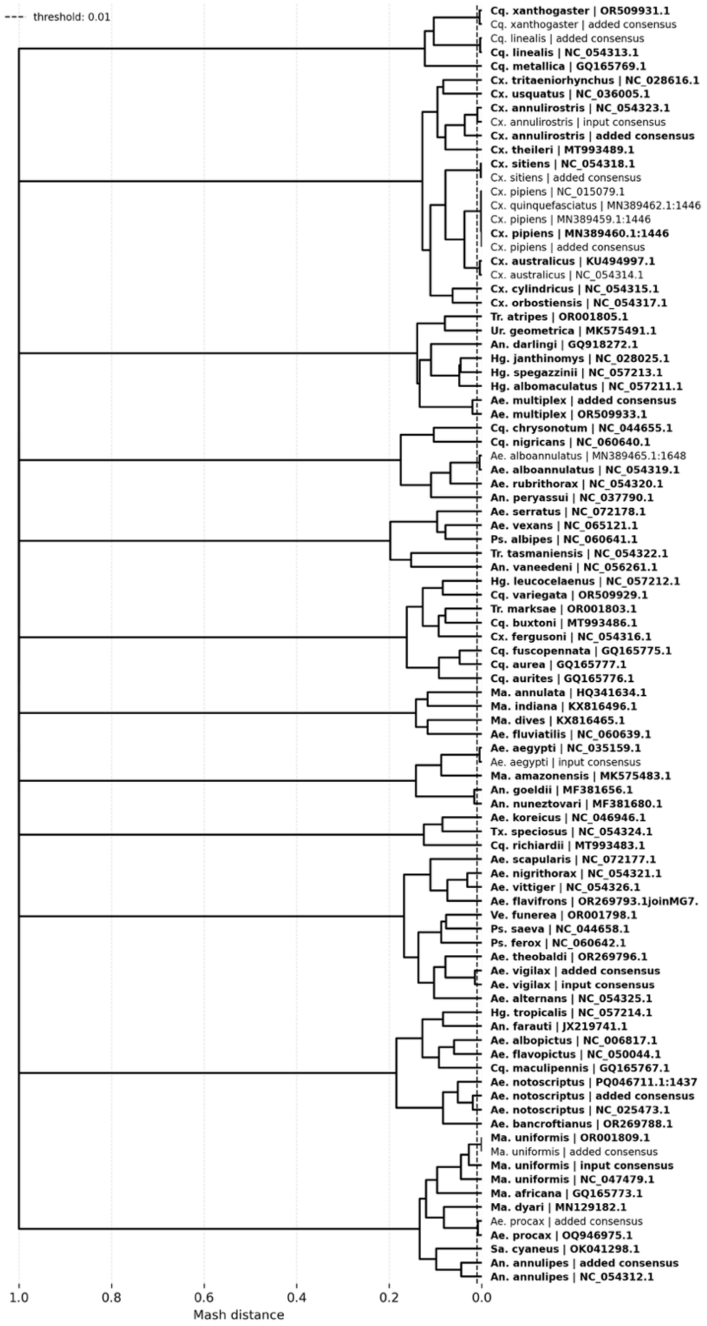


**Figure S12. Dendrograms showing Mash distances between sequences within the final COX1 species assignment reference set.** Dendrograms show clusters of pairwise Mash distances based on 31 bp k-mers between 931bp fragments of mosquito COX1 gene sequences used for species identification and relative abundance estimation. This does not represent a phylogenetic tree. Tips are annotated with species name and sequence identifier. Identifiers correspond to NCBI accessions for publicly available sequences, “input consensus” for consensus sequences generated from multiple published fragments, or “added consensus” for consensus sequences generated from available single-species pools, as described in Methods. Labels in bold indicate that sequence was retained in the final reference set. The dashed line indicates a Mash distance of 0.01 which was the threshold used to filter closely related references.

<u>Figure S13</u>

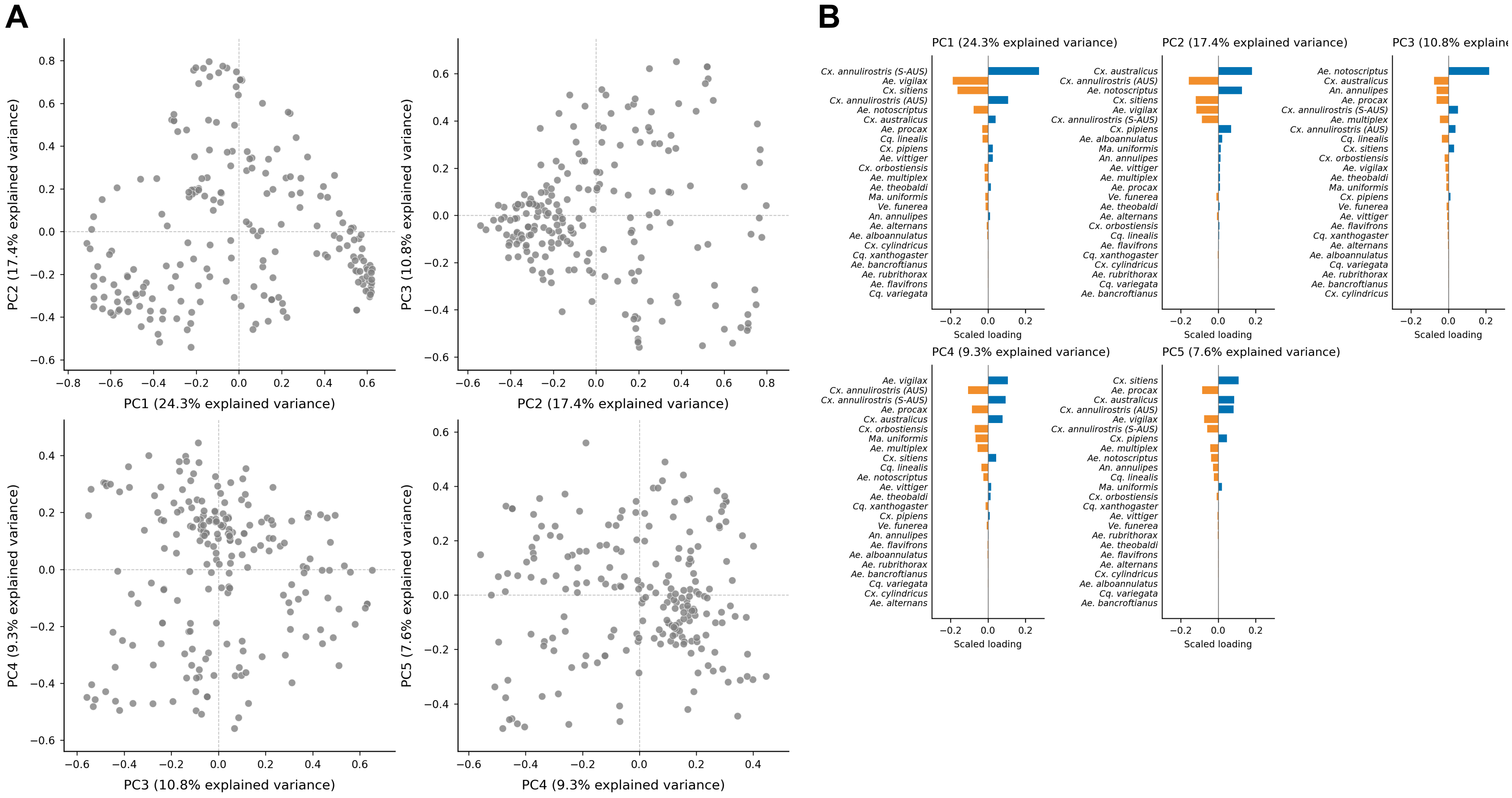


**Figure S13**. **Principal components of mosquito community composition across surveillance samples.** Principal component analysis was performed on Hellinger-transformed mosquito relative abundance estimates from 213 surveillance samples. **A)** Scatter plots show the distribution of samples across PC1 and PC2, PC2 and PC3, PC3 and PC4, and PC4 and PC5. Axis labels indicate the proportion of variance explained by each principal component. **B)** Barplots showing mosquito taxon loadings across PC1–PC5.

Figure S14

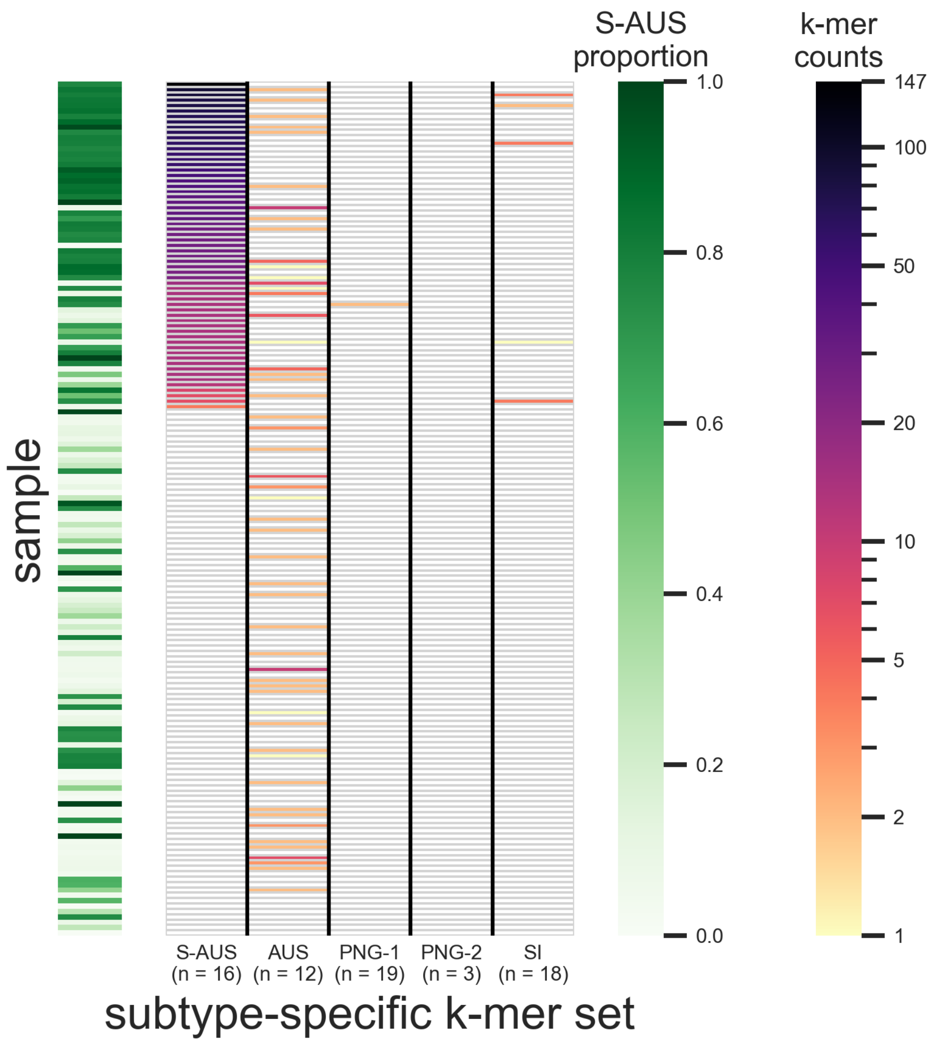


**Figure S14. Lineage-specific kmers for non-Australian *Culex annulirostris* populations are detected at low frequency within surveillance traps.** Previously published partial COX1 sequences from five *Culex annulirostris* mitochondrial lineages [30] were used to find 31 base pairs kmers that were present in at least 95% of individuals within the lineage, but no individuals outside of the lineage. Columns represent each lineage-specific kmer set and rows represent a surveillance trap with a positive *Cx. annulirostris* identification. The number of kmers within each set is written in the column label. Cell colour represents the summed raw count of kmers within each set identified in sequence data produced from the trap. On the left, a colour bar shows the estimated proportion of the S-AUS lineage within each trap, independently estimated using read pile-ups at lineage defining polymorphic sites.

Figure S15

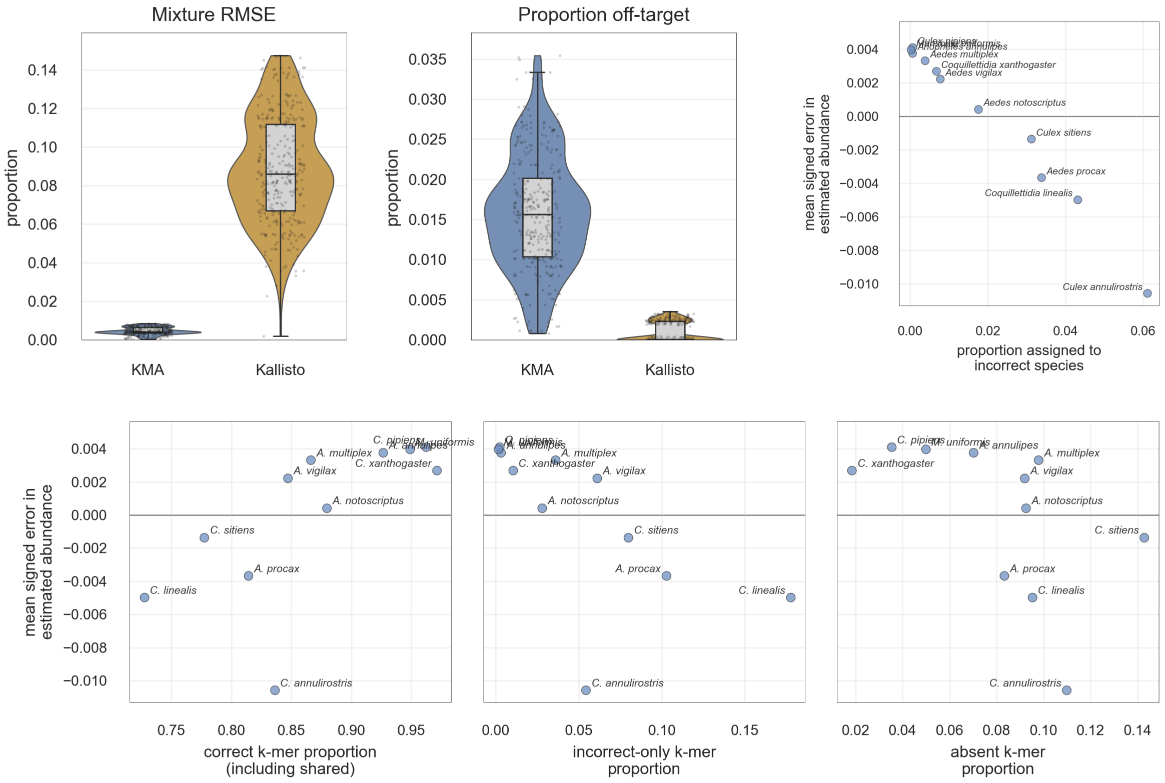


**Figure S15. Mosquito species identification and relative abundance estimation benchmarking.** Every four-species combination of 11 single species pools were generated in equal proportions (N=330). Prior to mixing, reads were aligned to three consensus sequences of COX1, soft clipped bases were removed, and reads were filtered to a minimum length of 100bp, as described in Methods. Reads in mixtures were then assigned to a reference set of COX1 sequences using KMA or Kallisto. Violin plots show **(A)** the root mean squared error (RMSE) computed from the estimated proportions compared to the expected proportions, and **(B)** the proportion of assigned reads mapped to a species not within the mixture. Results are separated by tool with KMA in blue and Kallisto in yellow. **(C)** Scatter plot showing the mean signed error in relative abundance estimation compared to 25% across mixtures (x-axis) against the proportion of reads from the corresponding single-species pool assigned to an incorrect species (y-axis). **(D)-(F**) Scatter plots showing mean signed error against k-mer representation metrics for each species: (D) the proportion of 31 bp k-mers shared between the sequence data and the correct reference sequence, including k-mers also shared with other species; (E) the proportion of 31 bp k-mers present only in incorrect reference sequences; and (F) the proportion of 31 bp k-mers absent from the reference set. Scatter plots were used to assess whether species-specific abundance-estimation error was associated with ambiguous or incomplete k-mer representation in the reference database.

**Supplementary Tables**

The following tables can be found within an accompanying excel spreadsheet.

Table S1
List of COSMOS probe targets

Table S2
Abundance estimates for mosquitoes across the surveillance samples

Table S3
Summary of virus detection results

Table S4
Mosquito COX1 reference panel sequences and sources

Table S5
Principal component temporal and spatial correlates association results

Table S6
S-AUS/AUS proportions in each trap and how grouped in Figure 4

**Supplementary Results**

Assessment of virus genome recovery in control samples

During the development of the presented workflow, we assessed the ability of COSMOS to recover viral genomes using an artificial construct containing eight concatenated fragments of seven arbovirus species' genomes, which produced read abundances proportional to their representation in the construct (**Fig. S3B**). In addition, COSMOS capture on a sample of Barmah Forest virus culture demonstrated complete genome recovery with 99.9% of the reference genome covered at a minimum depth of two sequence reads. Recovery by COSMOS tracked viral concentration across a dilution series: the number of sequenced unique virus reads was negatively correlated with PCR cycle threshold ($R^2$=0.999, N=4, P=0.0004) (**Fig. S3A**). This suggested COSMOS can recover near-complete viral genomes while preserving information about relative target abundance.

Batch specific assessment of index hopping

During assessment of virus coverage in our dataset of surveillance samples, we noticed that 20 samples carried detections of Barmah Forest virus (BFV) using our detection threshold of at least 5% of the genome covered at 2-fold depth. These samples had not been identified as carrying this virus via routine PCR testing performed by the NSW Arbovirus and mosquito surveillance program. They contained up to 25,851 BFV reads, producing up to 42% genome coverage. These samples were restricted to a single sequencing batch, COSMOS2, out of four batches processed during this study (COSMOS1-4). COSMOS2 also contained a sample of pure BFV virus culture which generated a large proportion of sequencing reads within the pool. Therefore, we suspected that, instead of representing a new detection, this signal may instead be a result of index hopping, whereby reads originating from one library are incorrectly assigned to another. To assess this, we compared virus recovery using Castanet with and without the use of the --PostFilt command line argument. –PostFilt removes reads with singleton start-end coordinate pairs, as these can arise through index hopping. The use of –PostFilt removed the BFV reads from the subset of samples within the COSMOS2 batch, leading us to suspect these were the results of index hopping (**Fig. S9)**.

The effect of using –PostFilt varied by batch. Virus coverage within COSMOS3 and COSMOS4 was highly similar with and without the use of –PostFilt, whereas COSMOS1 lost coverage across detections within a number of samples that were well-covered without –PostFilt (**Fig. S9**).

COSMOS1 was the first batch we performed COSMOS-capture on as part of this project, and it showed lower levels of read duplication compared to subsequent runs. This may have resulted in a greater proportion of non-index hopped singleton reads within this data which were removed by PostFilt, resulting in lower virus genome recovery in this batch with this use of --PostFilt (**Fig. S9).**

Based on these results, we applied the –PostFilt filter to COSMOS2 where there was direct evidence of index hopping, while –PostFilt was disabled for COSMOS1, COSMOS3 or COSMOS4. This batch specific approach removed apparent index-hopping signal while avoiding unnecessary loss of virus genome coverage across unaffected batches.

Recovery of the mosquito virome using metagenomic background in COSMOS sequencing data and association of *Culex australicus* with Wuhan Mosquito Virus 6

We set out to detect and quantify non-COSMOS targeted viruses in the surveillance samples (**Methods**). Sequences from these viruses are not enriched by the hybridisation capture step but are instead present in the library as background. Within a compiled reference set of 124 non-targeted viruses, 38 had more than 50% of their genomes covered, and coverage patterns showed apparent geographic structure when ordered west to east (**Fig. S2A**).

Whether these patterns are due to species or regional affinities requires further enquiry. We found that read counts recovered for Wuhan Mosquito Virus 6 (WuMV-6) strongly associated with the estimated relative abundance of *Culex australicus* across surveillance samples with mosquito abundance estimates ($R^2$=0.63, P=5.22 $\times 10^{-47}$, N=213), suggesting *Cx. australicus* may be an important host (**Fig. S2B**). WuMV-6 was originally discovered in China [35] and has since been suggested to have rapidly spread across the globe with evolutionary signatures suggestive of a cryptic vertebrate host [36]. Of 77 surveillance samples where WuMV-6 was detected, 44 produced at least 30% genome coverage with which phylogenetic analysis is possible, and 11 produced near-complete genomes (≥90% coverage). These results demonstrate that residual metagenomic sequence generated alongside COSMOS capture provides additional information, and further illustrates the ability of this workflow to investigate associations between viruses and mosquitoes.

Validation of a computational framework to estimate mosquito species relative abundance

Results presented in a pre-print by Forster *et al.* (2026)[19] have shown that Kallisto under-assigns reads when sample and reference sequences were separated by intraspecific diversity, which can lead to directional biases in relative abundance estimations. However, KMA assigns a greater proportion of reads to non-present references compared to Kallisto. Based on these findings, we decided to estimate relative abundance using counts of reads assigned by KMA but call a species as present only if identified by both Kallisto and KMA.

To develop our workflow and test our reference set, we used *in silico* mixtures of reads from the single species mosquito pools described above. Reads in these mixtures were assigned to our database using the tools KMA and Kallisto, and results from this test are shown in **Figure S5** and **Figure S15**. This includes a comparison of proportion estimation error between tools (shown as RMSE) and the proportion assigned to an incorrect species (**Fig. S15**). Our results corresponded to the simulation analysis by Forster *et al.*, which found that KMA was more accurate at proportion estimation but assigned a larger proportion of reads to incorrect species.

Across the mixtures, certain species were consistently slightly under-estimated and others over-estimated (**Fig. S5**). For example, *Culex annulirostris* was on average estimated as comprising 24% of the mixture, as opposed to the true mixture proportion of 25%. We found the level of error appeared to correspond to the proportion of reads assigned to an incorrect reference when the single species pool was assigned in isolation (**Fig. S15**). We assessed whether this was related to kmer sharing between the *Culex annulirostris* single species pool and other species references in the reference set, but did not observe unusually high sharing to other references, or an unusually low proportion of sharing with the *Cx. annulirostris* references compared to other pools.

One possible explanation for this is that the available references do not capture COX1 diversity within this species. *Cx. annulirostris* is known to comprise distinct mitochondrial COX1 lineages and intra-lineage genetic diversity within COX1 is also present [30]. To test this, we used CliqueSNV (v2.0.3) [37] to extract haplotypes from the *Cx. annulirostris* pool and added those with at least 5% frequency to the reference set (N=5). This increased the number of reads assigned to the reference set and produced more even estimates across mixtures (data not shown). This method is, by nature, highly dependent on database composition, and performance is likely to improve given greater coverage of genetic diversity across the sample set. We decided not to include CliqueSNV imputed references in the final reference set so as not to over fit to the validation samples, and because the results displayed in **Figure S5** support the overall accuracy of the final presented workflow.

Assessment of cryptic population structure in *Aedes notoscriptus*

Similarly to *Culex annulirostris*, multiple genetically distinct populations of *Ae. notoscriptus* have been identified across Australia [31]. Here, we clustered public sequences into three genetic populations: one more deeply divergent lineage (population 1) and two more closely related but distinct sister lineages (populations 2 and 3) (**Fig S8A**). We identified 15 population defining sites which differentiated population 1 from population 2 and 3, together with six sites that differentiated populations 2 and 3 and, as above, used these to estimate lineage proportions in surveillance traps.

Lineage proportions could be estimated for 97 out of 128 *Ae. notoscriptus* positive traps that had more than 20-fold coverage at 12 of the 15 population 1 informative sites and five of the six sites distinguishing populations 2 and 3 (**Fig. S8B**). The relative abundance of populations 2 and 3 varied along a north-south gradient, with population 2 more prevalent in northern NSW traps and population 3 in southern NSW (N=97, $R^2$=0.59, P<0.0001, computed using linear regression of latitude on proportion). In contrast, the proportion of population 1 did not appear to be associated with latitude (N=97, $R^2$=0.01, P>0.1, linear regression of latitude on population 1 vs. 2 or 3) (**Fig. S8B**). These patterns correspond to previous findings indicating the presence of northern and southern populations of *Ae. notoscriptus*, together with a third lineage occurring sympatrically across their ranges [31].
